# Longitudinal single cell transcriptomics reveals Krt8+ alveolar epithelial progenitors in lung regeneration

**DOI:** 10.1101/705244

**Authors:** Maximilian Strunz, Lukas M. Simon, Meshal Ansari, Laura F. Mattner, Ilias Angelidis, Christoph H. Mayr, Jaymin Kathiriya, Min Yee, Paulina Ogar, Arunima Sengupta, Igor Kukhtevich, Robert Schneider, Zhongming Zhao, Jens H.L. Neumann, Jürgen Behr, Carola Voss, Tobias Stöger, Mareike Lehmann, Melanie Königshoff, Gerald Burgstaller, Michael O’Reilly, Harold A. Chapman, Fabian J. Theis, Herbert B. Schiller

## Abstract

Lung injury activates quiescent stem and progenitor cells to regenerate alveolar structures. The sequence and coordination of transcriptional programs during this process has largely remained elusive. Using single cell RNA-seq, we first generated a whole-organ bird’s-eye view on cellular dynamics and cell-cell communication networks during mouse lung regeneration from ~30,000 cells at six timepoints. We discovered an injury-specific progenitor cell state characterized by Krt8 in flat epithelial cells covering alveolar surfaces. The number of these cells peaked during fibrogenesis in independent mouse models, as well as in human acute lung injury and fibrosis. Krt8+ alveolar progenitors featured a highly distinct connectome of receptor-ligand pairs with endothelial cells, fibroblasts, and macrophages. To ‘sky dive’ into epithelial differentiation dynamics, we sequenced >30,000 sorted epithelial cells at 18 timepoints and computationally derived cell state trajectories that were validated by lineage tracing genetic reporter mice. Airway stem cells within the club cell lineage and alveolar type-2 cells underwent transcriptional convergence onto the same Krt8+ progenitor cell state, which later resolved by terminal differentiation into alveolar type-1 cells. We derived distinct transcriptional regulators as key switch points in this process and show that induction of NFkB, p53, and hypoxia driven gene expression programs precede a Sox4, Ctnnb1, and Wwtr1 driven commitment towards alveolar type-1 cell fate. We show that epithelial cell plasticity can induce non-gradual transdifferentiation, involving intermediate progenitor cell states that may persist and promote disease if checkpoint signals for terminal differentiation are perturbed.

---

Lung disease is a major health burden accounting for one in six deaths globally^1^. The lung’s large surface area is exposed to a great variety of environmental and microbial insults causing injuries to its epithelium that require a regenerative response mediated by tissue-resident stem and progenitor cells. Epithelial lineage hierarchies in the mouse lung during development, adult homeostasis and injury have been studied using Cre recombinase-based genetic reporter systems. These studies revealed that depending on the location within the lung and the severity of injury, different stem and progenitor cell populations can be engaged^2–4^. Also, a surprisingly high degree of plasticity between epithelial cell type identities in injury scenarios has been noted^5^. The cell-intrinsic properties and niche signals driving these processes are not well understood and likely involve tight spatiotemporal control of crosstalk between the epithelium and the various immune and mesenchymal cell types that are activated or recruited after injury^6–8^.

In the alveolar compartment, gas exchange is enabled by ultra-thin extensions of alveolar type-1 pneumocytes (AT1) forming the alveolar surface area^9^. The surfactant-producing and cuboidal alveolar type-2 pneumocytes (AT2) have been shown to self-renew and act as progenitor cells by transdifferentiation into squamous AT1 cells, during both homeostatic turnover and injury^10^. In very severe cases of injury with massive loss of AT2 cells, both AT1 and AT2 cells can be replenished by airway-derived stem cell populations^11–15^. Recent evidence points towards the critical importance of inflammation-induced NFkappaB activation to drive AT2 cells into the cell cycle for alveolar regeneration^16^. However, the negative regulation of the NFkappaB pathway mediated by Yap/Taz in AT2 cells was also shown to be key for successful alveolar regeneration^17^. Moreover, the TGF-beta pathway has been proposed to mediate cell cycle arrest in AT2 cells followed by transdifferentiation into AT1 cells^18^. The molecular details and spatiotemporal organization of such decisive signals and pathways during recovery of the AT1 cell layer have not been resolved.

With the recent surge of single cell genomic methods, both experimental and computational, it is now possible to predict the future state of every single cell based on RNA velocity^19^, to model cell fate trajectories in pseudotime^20,21^, as well as infer the clonal history of single cells based on the allele frequencies of somatic mitochondrial mutations^22^, all within the same single-cell RNA sequencing (scRNAseq) experiment. These new methods are highly complementary with traditional lineage tracing in mice. High-resolution longitudinal single cell analysis of a dynamic system^23,24^, combined with new computational methods is unbiased and allows for discovery in high-throughput. Furthermore, the dynamics of cell-cell communication networks can be computationally approximated from scRNAseq data sets by the integration of receptor-ligand databases^23,25^. Here, we ask if we can leverage these ideas for the problem of gene regulation during epithelial regeneration. Recently, the first single cell studies of the mouse lung revealed the major cell type identities in normal homeostasis^26,27^ and aging^28^, and have led to the identification of an entirely new cell type, termed the pulmonary ionocyte^29,30^.

In this work, we charted the gene expression trajectories of 30 cell types in whole lung single cell suspensions after lung injury to provide a resource of the gene expression dynamics and routes of cell-cell communication during regeneration. We discover an intermediate alveolar epithelial progenitor cell state forming a unique cellular niche that peaks in frequency during the fibrogenic phase of tissue repair together with the appearance of myofibroblasts and M2-macrophages. Having surveyed whole lung dynamics, we employed a ‘sky dive’ approach^31^, by analyzing epithelial cell state transitions in a follow-up experiment with a very high temporal resolution for sorted epithelial cells. We discover transcriptional convergence of airway and alveolar cells onto Krt8+ alveolar progenitor cells and describe the sequence of transcriptional programs driving their commitment towards AT1 cell fate.

## Results

### A single cell portrait of lung regeneration

To comprehensively chart the cellular dynamics of all major cell lineages during regeneration after bleomycin-mediated acute lung injury, we collected whole-organ single cell suspensions from six timepoints after injury (day 3, 7, 10, 14, 21, and 28) and uninjured control lungs (PBS) with at least four replicate mice per timepoint. Using the Dropseq workflow^32^, we generated single cell transcriptomes from ~1000 cells per individual mouse, resulting in a final data set with 29,297 cells after quality control filtering (see Methods for preprocessing).

Single cell transcriptional profiles were visualized in two dimensions using the Uniform Manifold Approximation and Projection (UMAP) method^33^ and sampling timepoints were color-coded onto the UMAP embedding (Fig. 1a). Unsupervised clustering identified 46 distinct clusters, representing 28 cell type identities that were manually annotated using canonical marker genes and information from previously published scRNAseq data sets of the mouse lung^26,28^ (Fig. S1c, d; Table S1). We found good reproducibility of technical quality metrics and cell type assignment across the 28 mouse replicates (four per timepoint) (Fig. S1). Interestingly, this analysis revealed a gradual movement of cellular gene expression states along the regeneration time course, indicating that cell differentiation processes are at work (Fig. 1a). We observed good agreement between this scRNAseq data set with our previously published bulk RNAseq, as well as proteomics data from day 14 after bleomycin treatment^34^, validating the global injury-induced expression changes (Fig. 1b). Shared features of global bleomycin-induced changes in bulk transcriptomes, proteomes, and the scRNAseq data (Fig. 1c) were found to peak at day 10 and resolve during the regeneration time course (Fig. 1d). Interestingly, these shared features mainly showed cell type specific expression in the alveolar epithelium, fibroblasts and macrophages (Fig. S1g). To characterize the dynamic molecular changes within each cell type, we performed differential expression time course analysis (see Methods for details). A total of 6660 genes showed significant changes after injury in at least one cell type (FDR < 0.1, Table S2). The results of this analysis can be interactively explored with our webtool, which provides a user-friendly resource of gene expression changes in the whole lung during injury repair.

**Figure 1.**
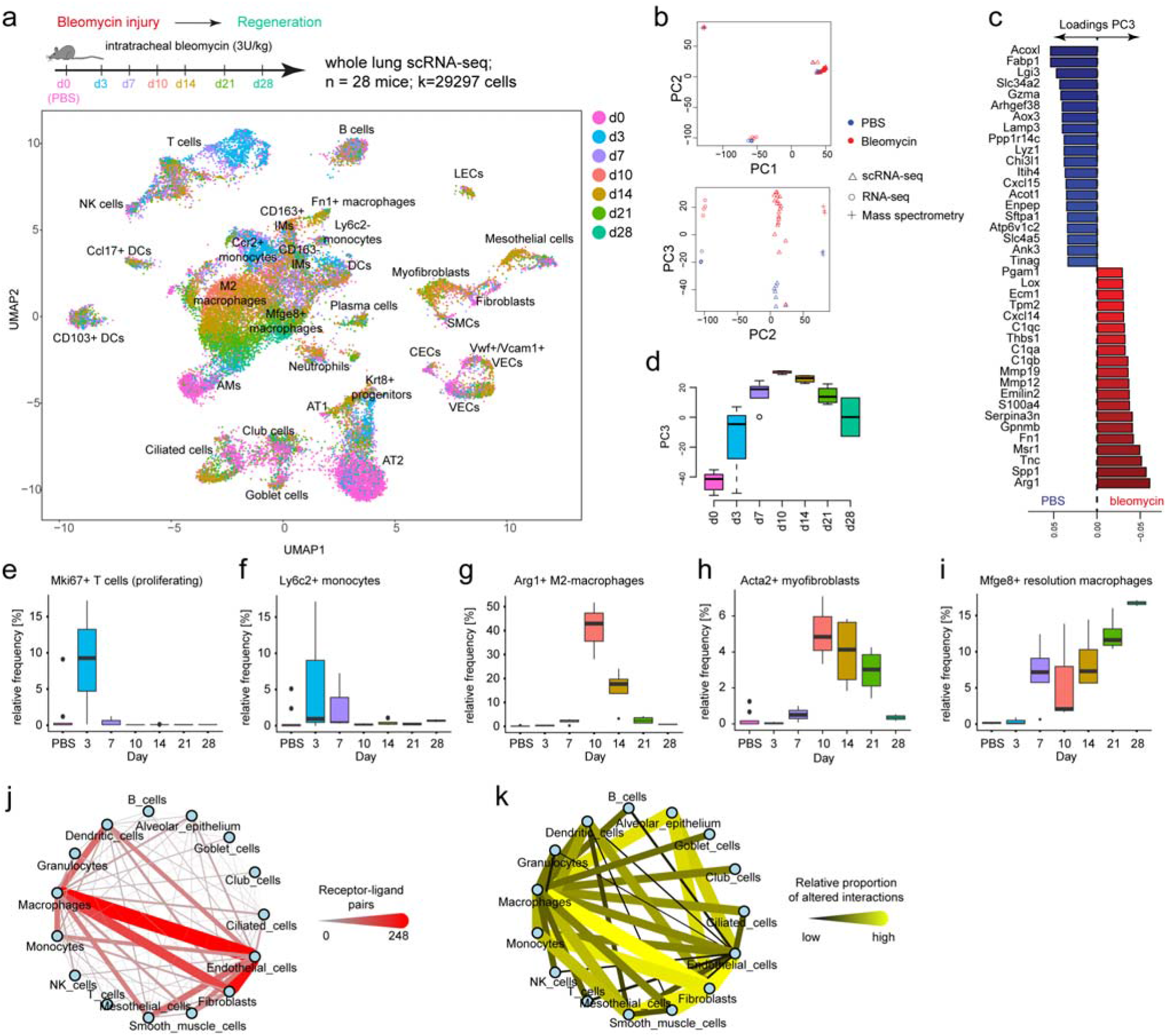
Longitudinal single cell RNA-seq analysis of lung regeneration reveals cell state and cell communication dynamics. (a) Single cell suspensions from whole mouse lungs were analyzed using scRNA-seq at the indicated time points after bleomycin-mediated lung injury. The color code in the UMAP embedding shows shifts of the indicated cell types in gene expression space during the regeneration time course. (b) Normalized bulk (RNA-seq) and *in silico* bulk (scRNA-seq) data were merged with proteome data (mass spectrometry) and quantile normalized. Bulk and protein data contain samples from day 14 after bleomycin-induced injury and controls^34^. The first two principal components show clustering by data modality. The third principal component separates bleomycin samples from controls across all three data modalities. Blue and red colors indicate control and bleomycin samples, respectively. (c) Barplot depicts genes with the highest loadings for principal component 3. (d) The box plot shows the time-resolved loading of PC3 peaking at day 10. The boxes represent the interquartile range, the horizontal line in the box is the median, and the whiskers represent 1.5 times the interquartile range. (e-i) Relative frequency of the indicated cell types relative to all other cells was calculated for individual mice at the indicated timepoints after injury (n=4) and for PBS treated control mice (n=7). The boxes represent the interquartile range, the horizontal line in the box is the median, and the whiskers represent 1.5 times the interquartile range. (j) The network shows 15 meta-cell type identities (see Fig. S1d) and their putative communication structure. Edge weight and color illustrate the number of receptor-ligand pairs between cell types. (k) The edges represent the relative proportion of receptor-ligand pairs between cell types with altered expression after injury.

Cell frequency dynamics showed an expansion of T cells early after injury at day 3 (Fig. 1e), recruitment of monocytes from blood within the first week after injury (Fig. 1f), the appearance of M2-macrophages peaking at day 10 (Fig. 1g), transient formation of myofibroblasts (Fig. 1h), and appearance of a resolution macrophage cell state peaking at day 28 (Fig. 1i). Myofibroblasts mediate fibrogenesis and have been shown to convert back to normal tissue-resident fibroblast phenotypes during lung fibrosis resolution^35^. In accordance with these findings, we observed a transient appearance of the myofibroblast cell state, which was marked by co-expression of *Col1a2*, *Acta2* and specific matrix genes such as *Tnc* (Fig. S2). Monocyte-derived macrophages have been shown to drive fibrogenesis^7,36,37^, at least in part via their interaction with fibroblasts^7^. It is still unclear how specific microenvironments shape the polarization of monocyte-derived macrophages into distinct subsets during injury repair. Previous experimental evidence suggests that in normal repair macrophages play a dual role in first promoting fibrogenesis and later promoting scar resolution. Subclustering macrophages revealed several distinct macrophage phenotypes at different timepoints after bleomycin injury (Fig. S3a-c). We used previously published bulk RNAseq signatures from lineage tracing of monocyte-derived macrophages in the bleomycin model^37^ to score individual cells, revealing that monocyte-derived macrophages give rise to both alveolar macrophages, as well as M2 and resolution macrophages (Fig. S3d, e), marked by *Arg1* and *Mfge8* (Fig. S3f, g) expression, respectively.

Next, we constructed a putative cell-cell communication network by mapping known receptor-ligand pairs across cell types (see Methods for details) (Fig. 1j). We integrated longitudinal dynamics by identifying receptor-ligand pairs where at least one of the genes showed a significant time course expression pattern in the corresponding cell type. The proportion of receptor-ligand pairs with altered expression over time was highest in possible communication routes between macrophages and fibroblasts, but also very striking differences in communication of these cell types with the alveolar epithelium were evident (Fig. 1k).

In summary, we have charted the gene expression kinetics over a four-week time course during lung regeneration for 28 individual cell types and predict altered niche environments and associated cell signaling during fibrogenesis that can be used for future spatial validations and functional studies.

### Alveolar regeneration involves a novel cell state

During annotation of cell type identity for the 46 clusters determined by the Louvain algorithm (Fig. S1c, d), we encountered a so far undescribed cell state in the alveolar epithelium, marked by expression of Keratin-8 (*Krt8*) and a highly distinct set of genes. Subclustering of alveolar epithelial cells resulted in four distinct clusters (Fig. 2a), which largely represented different timepoints (Fig. 2b). Notably, AT1 and AT2 cells were connected by cells mainly derived from intermediate timepoints, suggesting transdifferentiation dynamics. Hierarchical clustering of genes that were specific for these clusters revealed that two clusters contained AT2 cells marked by *Sftpc* expression, one representing the ground state and the other one an activated state marked by injury-induced genes, such as *Lcn2* and *Il33* (Fig. 2d, e). The cells in the unassigned cluster showed some similarity to AT1 cells, however, were clearly distinct and did not highly express the canonical marker genes for AT2 and AT1 (Fig. 2d, e). To analyze a possible transition of AT2 cells to these cells we used scVelo (see Methods for details) which uses the ratio of spliced to unspliced reads to infer RNA velocities^19^ and computationally predict the future state of individual cells. This RNA velocity analysis suggested that alveolar *Krt8*+ cells were derived from activated AT2 cells and might give rise to AT1 cells (Fig. 2c, d).

**Figure 2.**
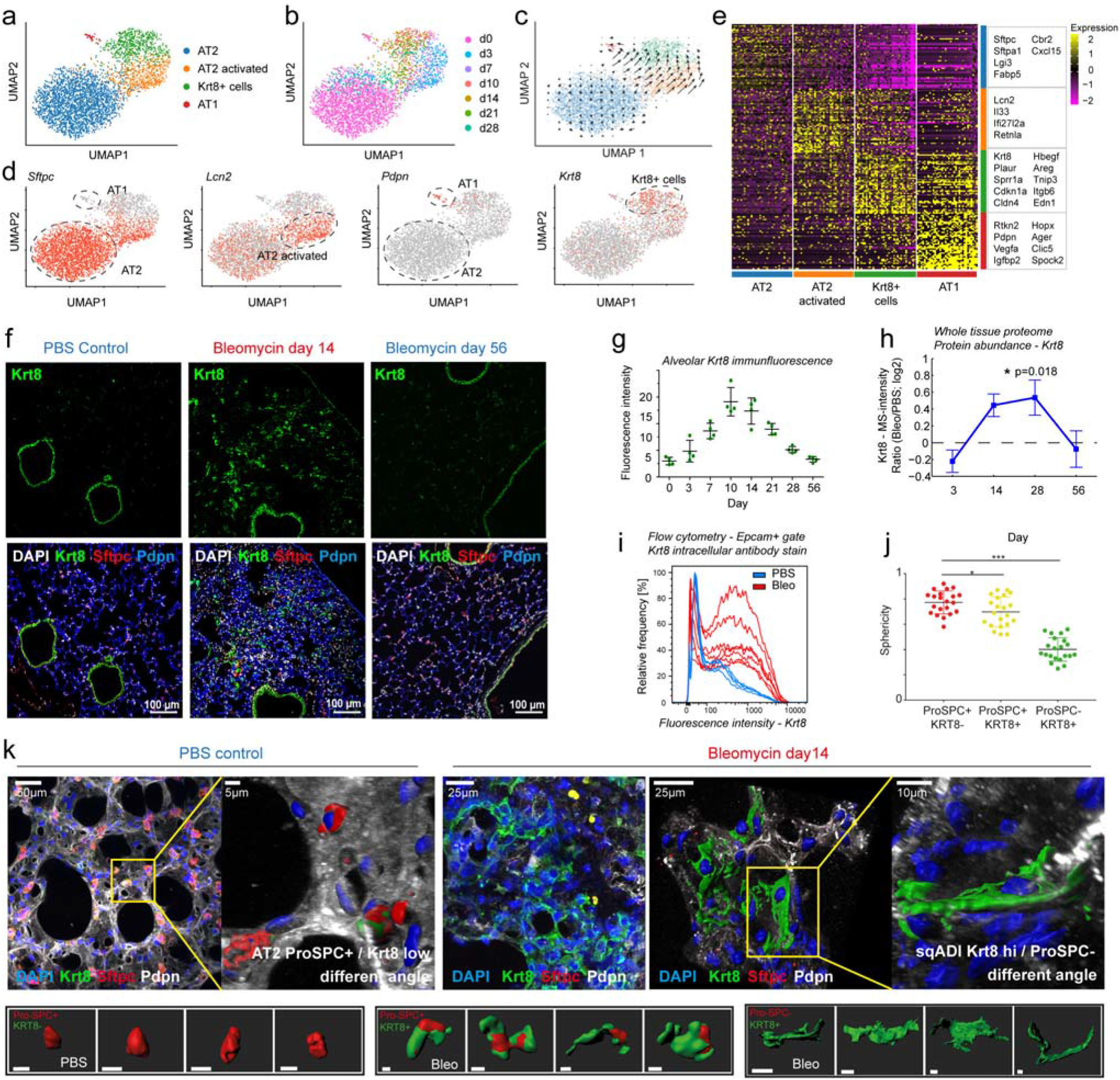
Alveolar regeneration features a transient squamous cell state marked by Krt8 expression. (a) UMAP embedding of alveolar epithelial cells shows four distinct gene expression states. (b) The color code shows the timepoints of sampling on the UMAP embedding. (c) RNA velocity information is plotted onto the UMAP embedding. Each arrow represents the local direction of transcriptome dynamics, estimated by comparing spliced vs. unspliced transcripts. Arrows are pointing towards the alveolar Krt8+ cell state after bleomycin-mediated injury. (d) UMAP embedded visualizations of single cells colored by gene expression for the four distinct gene expression states: AT2-*Sftpc*, AT2 activated-*Lcn2*, AT1-*Pdpn*, alveolar Krt8+-*Krt8*. (e) Heatmap shows the 50 most differentially expressed genes for the four alveolar cell states. The box shows gene names of selected examples. (f) Fluorescent immunostainings and confocal imaging of lung sections from the indicated conditions. Nuclei (DAPI) are colored in white, Krt8 appears in green, Sftpc (AT2 cells) in red, and Pdpn (AT1 cells) in blue. The scale bar indicates 100 microns. (g) Krt8 expression quantified by the mean fluorescence intensity of selected regions in the alveolar space, excluding Krt8+ airways. Alveolar Krt8 expression is highest at day 10 and day 14 (n=4 per timepoint, mean with SD). (h) Protein abundance of Krt8 in total lung homogenates was assessed by mass spectrometry^34^. The line plot shows the log2 ratio of Krt8 MS-intensity of mice at day 14 after bleomycin injury (n=4) and PBS control mice (n=4). Error bars show the standard error of the mean. (i) The histogram shows Krt8 fluorescence intensity quantified by flow cytometry using a CD45 negative and Epcam positive gate to select epithelial cells. PBS control mice (n = 5, blue color) and mice at day 10 after bleomycin (n=7, red color) are shown. (j) Alveolar cell sphericity analysis of 21 cells per condition revealed elongated cell shapes for Krt8+ cells in IF-stained precision cut lung slices (in k). Sphericity of 1 indicates round, cuboidal cells, 0 indicates flat cells. One-way ANOVA with Dunnett‘s post testing: *p=0.0376, ***p<0.0001. (k) Maximum projections of confocal z-stacks taken from immunostained 300 micron-thick precision cut lung slices (PCLS) are shown for a representative PBS control mouse and a mouse at day 14 after bleomycin injury. Nuclei (DAPI) are colored blue, Krt8 appears in green, Sftpc (AT2 cells) in red, and Pdpn (AT1 cells) in white. Small images below indicate the cell morphologies found in one healthy ROI within one PBS PCLS, and in two fibrotic ROI within one bleomycin PCLS. Note that Krt8+ cells form larger networks and clusters.

Immunostainings of Krt8 in lung sections confirmed its transient de novo expression in lung parenchyma from day 3 onwards. We observed a peak of *Krt8* expression in alveolar regions around day 10 - 14 after injury, which is also the peak of fibrogenesis (Fig. 2f, g; Fig. S4a). In contrast, the uninjured control lungs and fully regenerated lungs at eight weeks after injury showed *Krt8* expression only in airways (Fig. 2f). Interestingly, however, we occasionally found rare cells with high levels of *Krt8* expression in the alveolar space of uninjured control lungs (Figure S5), suggesting that the same cell state observed after injury may be a natural intermediate of homeostatic cell turnover. We also quantified the transient burst of Krt8 protein expression in injured alveolar space using mass spectrometry-based whole tissue proteomics (Fig. 2h) and flow cytometry (Fig. 2i; Fig. S4b, c). On the gene expression level, we found striking differences between alveolar Krt8+ cells and the related AT1 cells, including cornifin-A (*Sprr1a*), which interestingly has a known function in formation of the cornified envelope in keratinocytes and thus may have a function in squamous differentiation of lung cells. Alveolar Krt8+ cells also featured high expression of the low-affinity epidermal growth factor receptor ligands *Areg* and *Hbegf*, as well as the integrin *Itgb6*. Flow cytometry confirmed the increased expression of Itgb6 on lung epithelium after injury and the high expression of Itgb6 on the surface of alveolar Krt8+ cells (Fig. S4b, c). We also validated the increased expression of Areg and Hbegf in lung sections after bleomycin injury and observed partial co-localization with alveolar Krt8+ cells (Fig. S4d, e).

To analyze the morphology of alveolar Krt8+ cells, we fixed 300 micron-thick precision cut lung slices that encompass the whole alveolus for immunostainings and used confocal microscopy for morphometric analysis (Fig. 2k). While Krt8 is expressed only at very low levels in cuboidal AT2 cells in the uninjured lung, we observed vastly increased expression in still cuboidal AT2 cells expressing Sftpc at day 14 after injury and a large number of alveolar Krt8+ cells with no Sftpc expression and squamous morphology. In comparison to AT2 cells, the Krt8+ cells showed a significantly reduced sphericity factor and also AT2 cells with upregulated Krt8 after injury were found to assume a significantly flatter shape (Fig. 2j). Thus, we conclude that the Krt8+ cells mostly assume a flat squamous shape and cover the alveolar surface similar to AT1 cells.

To determine if the appearance of alveolar Krt8+ cells is specific to the bleomycin injury model, we turned to two other independent mouse models that are not based on DNA damage for the injury. The neonatal exposure of mice to hypoxic or hyperoxic conditions leads to higher susceptibility of these mice to injury and fibrosis upon adult infection with an Influenza type-A strain^38–40^. We assessed the expression of Krt8 in the alveolar space after influenza infection and observed high numbers of Krt8+ cells in both the neonatal hypoxia and hyperoxia cohort, but not in their littermate controls that were exposed to normal room air after birth, and thus did not contract a major injury after infection (Fig. S6a). Exposure of adult mice to hyperoxia has been shown to preferentially kill alveolar AT1 cells^41^. We further observed the appearance of alveolar Krt8+ cells in mice already at day 3 after a 60-hour exposure to hyperoxia (Fig. S6b).

To test if KRT8+ alveolar cells can also be observed in human acute lung injury and chronic lung disease associated with alveolar injury, we stained human tissue sections and did not detect any expression of KRT8 in the alveolar space of non-injured control lungs (n=7; Fig. 3a). In sharp contrast, we observed very strong alveolar KRT8 expression in human acute respiratory distress syndrome (ARDS, n=2) caused by Influenza-A and pneumococcal infection and interstitial lung disease patients with various diagnoses (n=5; Fig. 3b, c).

**Figure 3.**
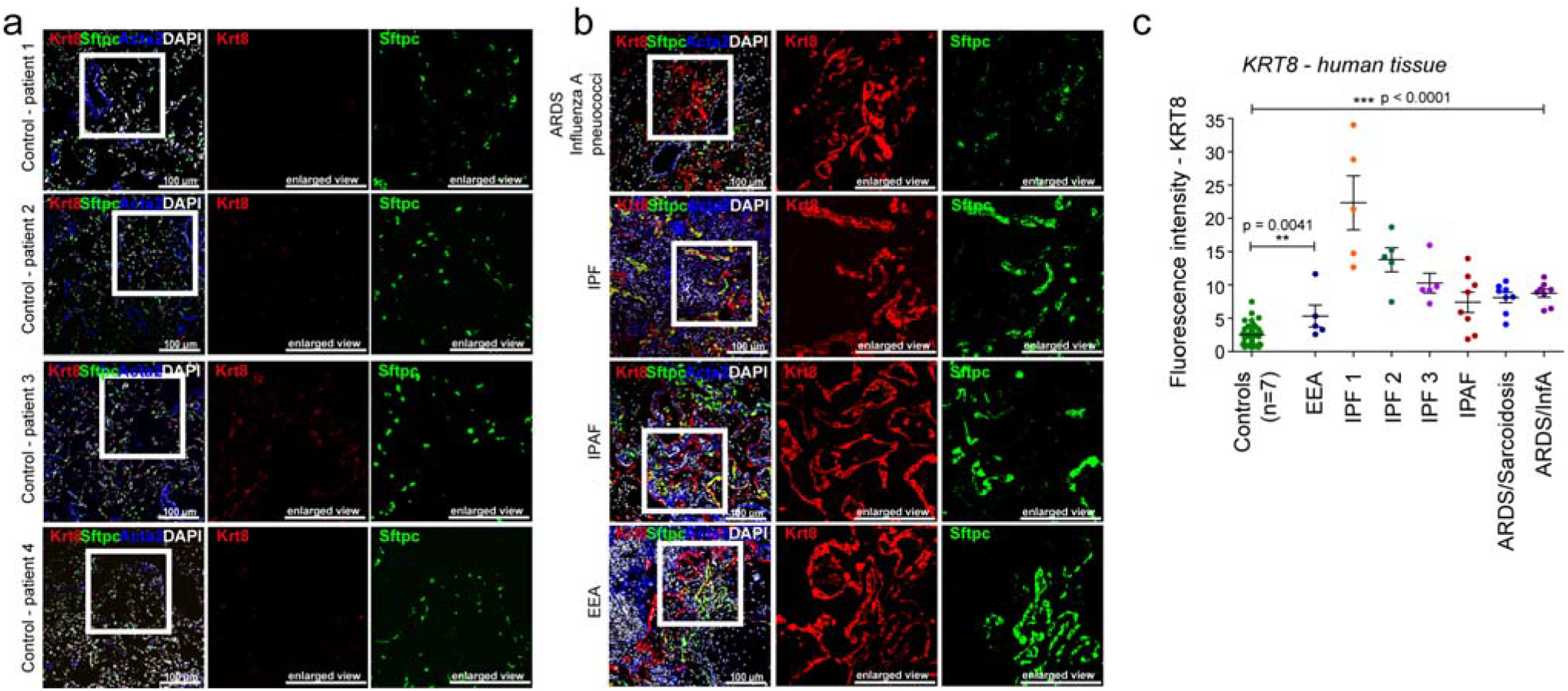
Krt8+ alveolar cells appear in human acute lung injury and fibrosis. (a) FFPE sections from non-fibrotic control parenchyma derived from non-involved areas in tumor resections were stained against Krt8 (red), Sftpc (green), and Acta2 (blue). Scale bar = 100 microns. (b) Human lung tissue sections were stained as in (a) revealing pronounced Krt8 expression at the site of acutely injured lesions (ARDS diagnosis) and fibrotic regions of ILD patient lungs (IPAF, IPF and EAA diagnosis). Scale bar = 100 microns. (c) Fluorescence intensity of Krt8 stainings was quantified from 5-8 representative areas of control tissue (n=7), ILD tissue (n=5), and ARDS (n=2). One-way ANOVA statistical analysis: ***p<0.0001, **p=0.0041.

In summary, we have discovered a highly distinct squamous epithelial cell state in alveoli during lung regeneration that appears transiently during the fibrogenic phase after injury. We hypothesized that the alveolar Krt8+ cells are progenitor cells derived from AT2 cells that later resolve into mature AT1 cells.

### A sky dive into epithelial cell fate transitions

The whole lung survey showed an appearance of a novel alveolar cell state peaking during the fibrogenic phase of tissue repair around day 10. We used these results for the experimental design of a second follow-up experiment with higher temporal and cellular resolution that aimed at modeling the generation of alveolar Krt8+ cells from epithelial progenitor populations. Sorting EpCam+ cells before single cell sequencing enabled us to focus our analysis on the epithelial cell compartment. To obtain high resolution for the cell state trajectory towards Krt8+ cells, we performed daily sampling of single cell transcriptomes up to day 13. We also included later timepoints up to day 54 after injury to analyze the recovery of the system back to the baseline with fully regenerated AT1 cells. In total, we collected 18 timepoints after injury using two replicate mice each (n=36 mice; k=34575 cells) (Fig. 4a).

**Figure 4.**
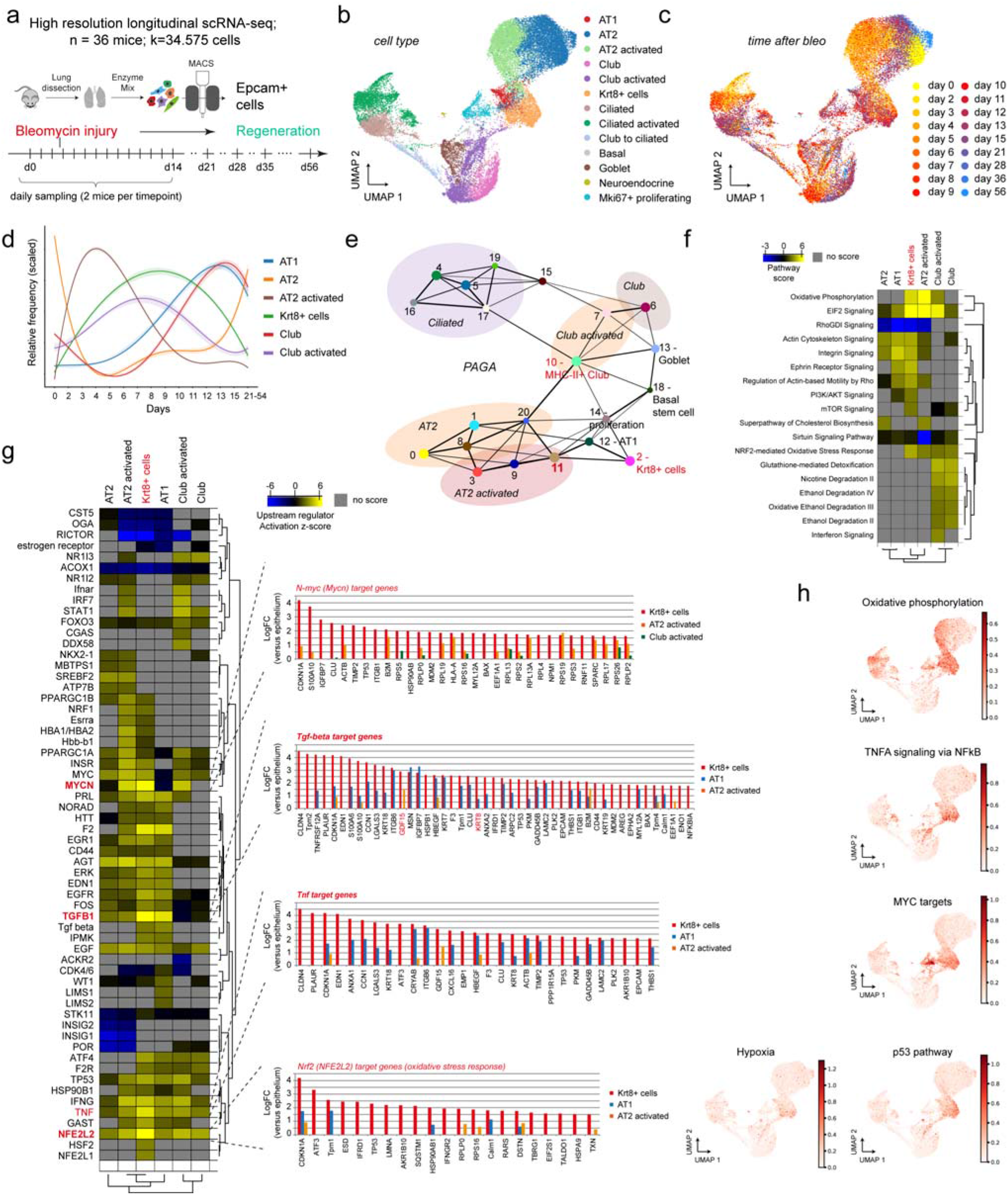
High-resolution sampling of epithelial cell state transitions reveals intermediate pathways and transdifferentiation bridges. (a) A high-resolution longitudinal data set was generated by subjecting sorted cells from the epithelial compartment to scRNAseq from the 18 indicated timepoints. UMAP embedding displays cells colored by cell type identity (b) and timepoint (c). (d) The lines represent smoothed relative frequencies of alveolar cell types and club cells over the time course. Confidence interval derived from the smoothing fit is shown. (e) PAGA graph visualizes potential cell-type transitions and the topology of the data manifold. Nodes represent Louvain clusters and thicker edges indicate stronger connectedness between clusters. (f) Ingenuity pathway analysis was used to score the activity of pathways within the signatures of the indicated cell states. The pathway activation z-scores were grouped by hierarchical clustering using their Pearson correlation. (g) Ingenuity upstream regulator analysis was used to score the activity of upstream regulators within the signatures of the indicated cell states. The activation z-scores were grouped by hierarchical clustering using their Pearson correlation. Bar graphs show target genes sorted by highest expression in Krt8+ cells relative to all other cells. (h) UMAP embedded visualizations of single cells colored by gene expression signature scores for the indicated pathways.

Cell type identities in the high-resolution epithelial cell data set were consistent with the first experiment. However, in addition, we identified neuroendocrine cells and basal stem cells that were not observed in the whole lung survey (Fig. 4b; Fig. S7). Some cell types exhibited remarkable heterogeneity in their transcriptional states along the tissue regeneration time course, so that we annotated some of the clusters as ‘activated’ cell states early after injury (Fig. 4b, c; Table S3). The increased relative frequency of activated AT2 cells preceded the transient appearance of Krt8+ cells, while the frequency increase of activated airway club cells was observed later than the activated AT2 cells (Fig. 4d). To analyze global connectivity and potential trajectory topology we applied partition-based graph abstraction (PAGA), which provides an interpretable graph-like map of the data manifold^42^ (Fig. 4e) and has been scored as one of the top performing pseudotime methods in a recent benchmark^20^.

Interestingly, the PAGA map revealed several nodes with high connectivity between cell types that represented potential transdifferentiation bridges. In particular, we observed a subset of airway club cells (cluster 10) with strong connectivity to the alveolar cells including Krt8+ cells, and an activated AT2 cell state (cluster 11) which also featured high connectivity to Krt8+ cells.

Differential gene expression analysis between the club cell subsets revealed that two clusters (6 and 7) mainly represented club cells at different times after injury, which we termed ‘club’ and ‘club activated’, respectively. Cluster 10, however, was highly distinct and most prominently marked by high expression of MHC-II complex genes (e.g. *H2-Ab1*) and the cysteine proteinase inhibitor Cystatin-C (*Cst3*), which is typically co-expressed with MHC-II in dendritic cells^43^ (Fig. S8a-d). Activated club cells were marked by the upregulation of ribosomal genes and ATP synthesis, and downregulation of the Wnt signaling pathway (Fig. S8e). The MHC-II+ club cell subset also featured higher ribosomal content, cytokine production, and expression of intermediate filaments, such as Vimentin. Using immunofluorescence, we validated the selective expression of *Cst3* in a subset of Scgb1a1+ airway club cells (Fig. S8f), suggesting that increased MHC-II complex expression could be a dynamic response to injury within the airway epithelium.

Activated AT2 and club cells, as well as the Krt8+ cells, were characterized by increased oxidative phosphorylation (Fig. 4f). This metabolic switch is often observed in cellular differentiation processes^44^. The activated cell states and Krt8+ cells were also characterized by increased ribosomal content, ElF2 signaling components, and N-Myc target genes, suggesting that these cells had re-entered the cell cycle (Fig. 4f, g). Both the activated AT2 cells and the MHC-II+ club cell subset connected with a cluster of proliferative cells (cluster 14) marked by *Mki67* and *Top2a* expression (Fig. 4e). The relative frequency of these proliferating cells peaked at day 15 (Fig. S9a). Counting Ki67+ cells in immunostainings confirmed the peak of cell proliferation around day 14 with a sudden drop in proliferation rates around day 28 (Fig. S9d, e). We performed cell cycle regression within the proliferative cells to deconvolve cell type identity (Fig. S9b), revealing that Krt8+ cells, AT2, club, and the MHC-II+ club cells all proliferated after injury (Fig. S9c). We validated proliferating Krt8+ cells in co-immunostainings, which indeed showed Ki67+ nuclei of Krt8+ cells at day 10 after injury (Fig. S9f). In comparison to the other epithelial cells, Krt8+ cells were significantly enriched for TNF, TGF-beta and NRF2 pathway (hypoxia response) target genes (Fig. 4g). Indeed, scoring the hypoxia response, p53, MYC, TNFA, and oxidative phosphorylation pathways onto the UMAP embedding showed a specific enrichment of pathway genes on the Krt8+ cluster (Fig. 4h).

### Transcriptional convergence of AT2 and club cells towards Krt8+ cells

RNA velocity vectors overlaid onto the UMAP embedding predicted transdifferentiation of club cells towards ciliated and goblet cells, which is in agreement with previous literature^2^ (Fig. 5a). Interestingly, RNA velocities also strongly suggested a dual origin of alveolar Krt8+ cells from AT2 and airway club cells (Fig. 5a, b). We restricted the analysis to these three groups and calculated terminal state likelihoods based on RNA velocities, which suggested differentiation of activated AT2 and airway club cells towards a Krt8+ cell endpoint (Fig. 5b).

**Figure 5.**
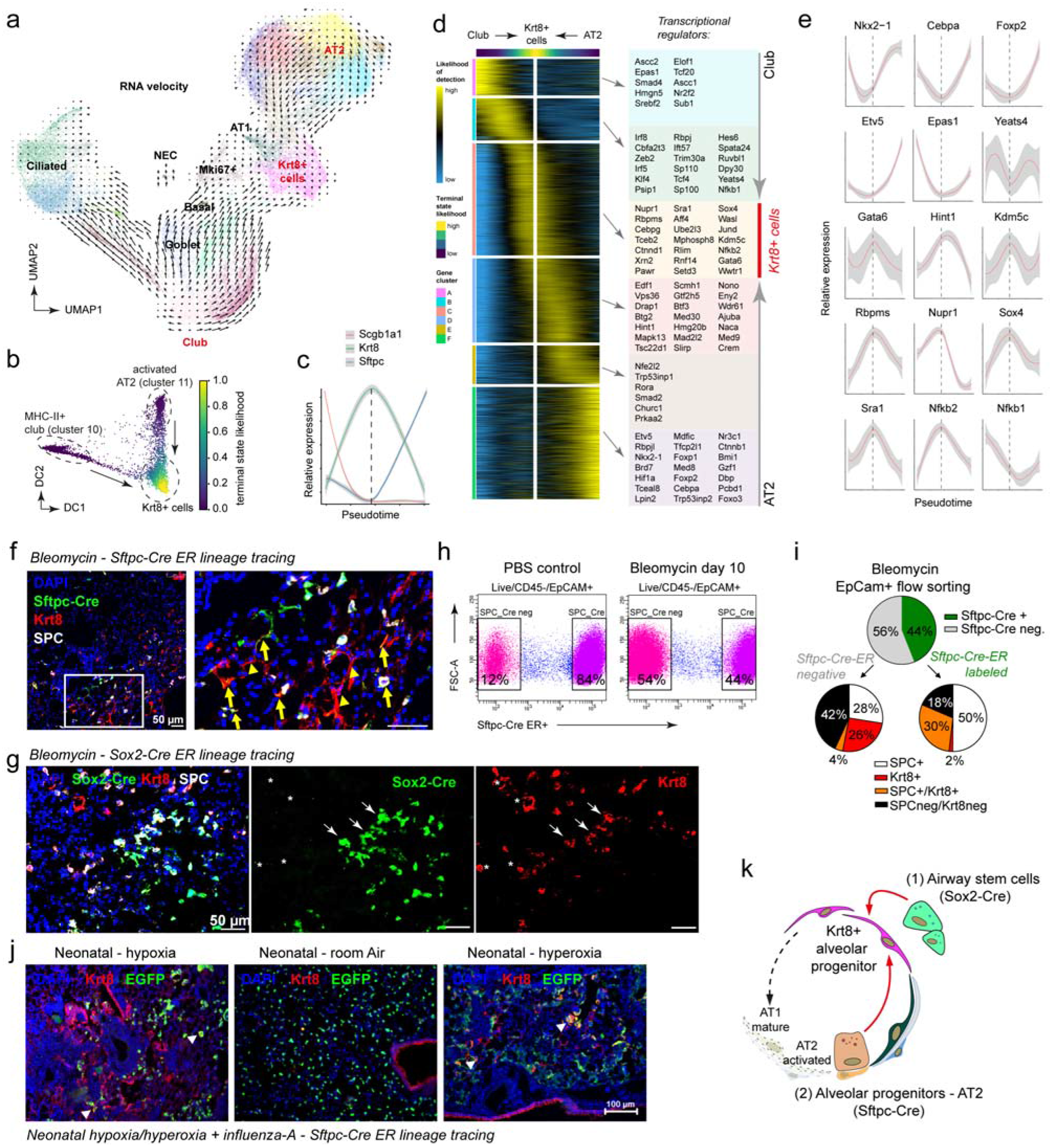
Transcriptional convergence of club and AT2 cells onto the alveolar Krt8+ cell state. (a) Velocity plot displays the UMAP embedding colored by Louvain clusters with velocity information overlaid (arrows), indicating convergence of AT2 and club cells onto the alveolar Krt8+ cell state. (b) Diffusion map of Louvain clusters 2, 10, and 11 colored by inferred terminal state likelihood reveals two distinct transdifferentiation trajectories from activated AT2 and MHC-II+ club cells towards a Krt8+ cell state. (c) The lines illustrate smoothed expression levels of *Scgb1a1, Krt8*, and *Sftpc* across the trajectory, marking cell identities. The dashed vertical line indicates the peak of *Krt8* expression. Grey colors represent the confidence interval derived from smoothing fit. (d) The heatmap shows the gene expression patterns along the differentiation trajectory based on the inferred likelihood of detection for 3036 altered genes. (e) Line plots show the smoothed relative expression levels of selected transcriptional regulators across the converging trajectories. The dashed vertical line indicates the peak of *Krt8* expression. Grey colors represent the confidence interval derived from smoothing fit. (f) Both Sftpc-Cre/ER-labeled and Sftpc-Cre/ER-negative Krt8+ cells are found in the lungs of mice at day 10 post bleomycin injury. Scale bar = 50 microns. Arrowheads show mostly squamous looking Sftpc-Cre/ER-negative Krt8+ cells, while arrows identify Sftpc-Cre/ER-labeled SPC+/Krt8+ cells. (g) Sox2-Cre lineage trace identifies airway derived (Sox2-labeled) Krt8+ cells. (h) Flow cytometry plots of single cell suspensions prepared from either saline (n=2) or bleomycin treated (n=2) mice. While ~12% of the EpCam+ cells are Sftpc-Cre negative (airway cells), they account for ~54% of total epithelial cells 10 days after bleomycin injury. (i) Quantification of total epithelial Krt8+ cells (EpCam+/Krt8+) revealed that 44% of the Krt8+ cells in day 10 bleomycin injured lungs are Sftpc-Cre/ER-labeled (AT2 derived) and 56% of the epithelial Krt8+ cells are Sftpc-Cre/ER-negative, and therefore most likely coming from an airway lineage. Further analysis revealed that the majority of the SPC+/Krt8+ cells are Sftpc-Cre ER-labeled (30% of total traced cells). n=2 mice used for differential quantification on a cytospin. At least 800 cells were counted for each condition. (j) Lungs of infected Sftpc^CreERT2^; Rosa26R^mTmG^ mice show EGFP expression in AT2 cells (green) and were stained for Krt8 (red). Scale bar = 100 microns. (k) Lineage tracing experiments and scRNAseq experiments show that both airway progenitors and AT2 cells give rise to the same Krt8+ cell transcriptional state.

To understand the transcriptional dynamics during this process we performed pseudotemporal trajectory analysis and identified 3036 genes showing distinct expression patterns along the trajectories (Fig. 5c, d; Table S4). We observed a gradual decline in expression of the Homeobox protein *Nkx-2.1*, critical for lung development and lung epithelial identity^45^, as well as *Foxp2*, which is one of the key transcriptional repressors involved in the specification and differentiation of the lung epithelium^46,47^, in both club and AT2 cells during conversion to Krt8+ cells (Fig. 5e). Also, expression of the transcription factor *Cebpa* with important functions in lung development and maintenance of both club and AT2 cell identity^48–50^ reached a minimum at the Krt8+ state. AT2 cell conversion into Krt8+ cells was marked by a drastic reduction of the transcription factor *Etv5*, which has been shown to be essential for the maintenance of AT2 cells^51^ (Fig. 5e). Conversely, the differentiation towards the Krt8+ cell signature expression was characterized by a gradual increase in one of the master regulators of AT1 cell differentiation *Gata6^52,53^* in both club and AT2 cells. Both trajectories converged on a large number of alveolar Krt8+ cell specific genes representing distinct pathways (Fig. 4) and their transcriptional regulators, including the stress-induced p53 interactor *Nupr1*, the master regulator of epithelial to mesenchymal transition *Sox4*, and many other genes including chromatin remodeling factors such as the histone demethylase *Kdm5c* (Fig. 5e).

In order to validate the predictions from this computational analysis, we used the Sftpc-Cre (AT2 cells) and Sox2-Cre (airway cells) reporter mice to lineage trace the origin of Krt8+ cells. Both Sftpc-Cre/ER-labeled and Sftpc-Cre/ER-negative Krt8+ cells were found in the lungs of mice post bleomycin injury, indicating that Krt8+ cells were indeed partially airway-derived (Fig. 5f). Flow cytometry analysis showed that while in single cell suspensions isolated from control lungs we found 84% of EpCam+ cells to be AT2-derived, this was reduced to 44% after bleomycin injury (Fig. 5h). To better quantify the proportion of AT2-derived Krt8+ cells, we flow sorted EpCam+ cells and evaluated 800 single cells on cytospins for expression of *Krt8*, *Sftpc* and the lineage label. We found that in this analysis 26% of lineage negative cells were *Krt8*+, while most AT2-derived cells with high levels of *Krt8* still expressed *Sftpc* (Fig. 5i). To prove that *Sftpc*-Cre lineage negative Krt8+ cells were airway-derived, we turned to the *Sox2*-Cre reporter mice, which specifically report the airway lineage and again found that only a fraction of the Krt8+ cells was *Sox2* lineage labeled (Fig. 5g). Similar to this analysis, we also found a dual origin of Krt8+ cells in the neonatal hyper-/hypoxia with adult influenza model. Keratin-8 stainings in lung tissue from Sftpc-Cre reporter mice after influenza infection revealed that only some of the Krt8+ cells were AT2-derived (Fig. 5j).

Together, these results show that both airway-derived progenitors within the club cell lineage and AT2 cells converge on the same transcriptional state during regeneration of alveoli (Fig. 5k). The data reveals the transcriptional regulators of this process and highlights the dynamic interplay of several inflammation- and stress-induced transcriptional regulators with master regulators of lung epithelial identity such as *Cebpa*, *Nkx2.1*, *Foxp2,* and *Etv5*.

### Krt8+ cells are alveolar progenitors that reconstitute the AT1 layer

We have shown that the Krt8+ cell state is derived from airway progenitors and AT2 cells and that it appears transiently, resolving between day 14 and day 54 after injury in the bleomycin model. Thus, our hypothesis was that these cells are alveolar progenitors that undergo terminal differentiation into mature AT1 cells during this time. To investigate and model this potential cell trajectory, we reanalyzed the subset of Krt8+ cells and AT1 cells separately. Indeed, velocity information overlaid on to the derived UMAP embedding indicated differentiation from Krt8+ cells towards AT1 cells (Fig. 6a). Moreover, analysis of the ratio of spliced and unspliced reads revealed gradual induction of transcription of the AT1 cell marker *Ager* in cells during the fibrogenic phase around day 14 (Fig. 6b). Days 0, 36 and 56 contained a significantly lower ratio of unspliced over spliced *Ager* reads compared to all other timepoints (Fig. 6c; Wilcoxon Rank Sum test, P<1e-46), marking the steady-state ratio of mature AT1 cells. This mRNA velocity information predicted a gradual increase of *Ager* expression along the cell trajectory towards the AT1 terminal differentiation state (Fig. 6d), which was correspondingly reflected by the observed total expression levels of *Ager* (Fig. 6e). Using this information, we calculated a pseudotime trajectory and determined gene expression dynamics for 1150 significantly regulated genes along the Krt8+ alveolar progenitor to AT1 transition (Fig. 6f; Table S5).

**Figure 6.**
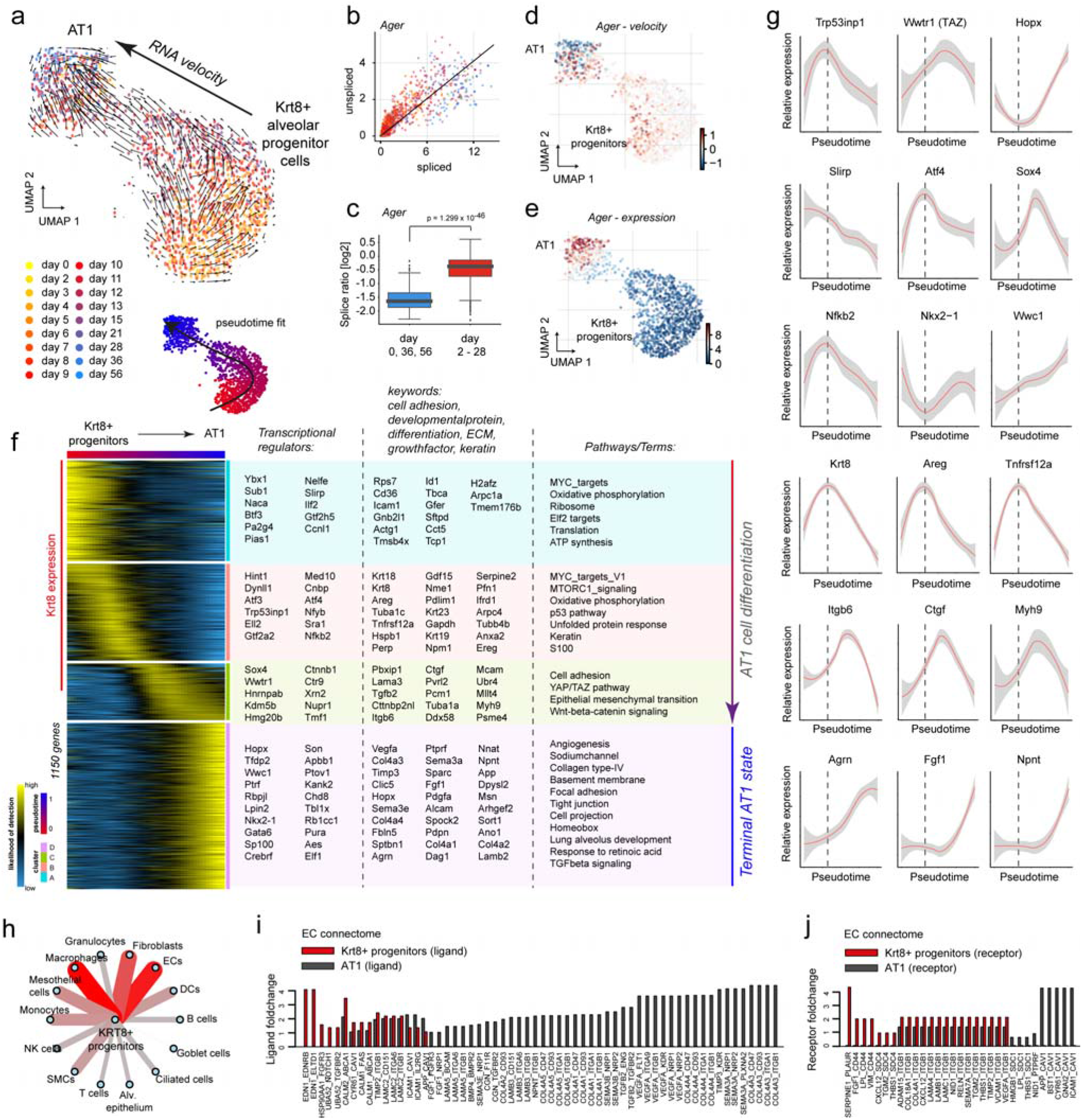
Pseudotime analysis reveals Krt8+ cells as alveolar progenitors that give rise to AT1 cells. (a) Velocity plot displays the UMAP embedding colored by timepoint with velocity information overlaid (arrows), indicating terminal differentiation of Krt8+ progenitors into AT1 cells. (b) Velocity phase plot shows the number of spliced and unspliced reads of the AT1 marker *Ager* for each cell (points) on the X and Y axes, respectively. Cells are colored by timepoint and the black line represents the linear steady-state fit. Cells above and below the diagonal are predicted to be in inductive or repressive states, respectively. (c) Boxplot shows the log2 ratio of unspliced over spliced *Ager* reads for days 0, 36 and 56 (blue) and all other timepoints (red). To avoid division by zero, one was added to both counts. UMAP embedding colored by *Ager* velocity (d) and expression (e) displays a gradual increase along the inferred trajectory. (f) The heatmap shows the gene expression patterns across the differentiation trajectory for 1150 altered genes. (g) The line plots illustrate smoothed expression across the differentiation trajectory for a number of exemplary genes. Grey colors represent the confidence interval derived from the smoothing fit. The dotted line indicates the peak of Krt8 expression. (h) The cell-cell communication network displays the number of receptor-ligand pairs between the molecular markers of the Krt8+ alveolar progenitor cell state and all other meta cell type identities (Figure 1). (i, j) The bar graphs show the average log2 fold change of either receptors (i) or ligands (j) within the endothelial cell (EC) connectome for Krt8+ alveolar progenitors and AT1 cells.

The pseudotime trajectory revealed distinct phases within the Krt8+ alveolar progenitor cell population that we modeled as a gradual maturation towards the mature AT1 cell state. Thus, heterogeneity within the Krt8+ alveolar progenitor cells (cluster 2) can be explained by time. We split the transdifferentiation trajectory into 4 phases that were marked by distinct sets of transcriptional regulators, developmental genes and signaling pathways (Fig. 6f, g). The initial phase was marked by genes and pathways consistent with cell growth after exiting the cell cycle (e.g. MYC targets). This was accompanied by the induction of stress-related signaling pathways, such as the p53 pathway and the unfolded protein response pathway, featuring increased expression of the corresponding transcriptional regulators such as *Trp53inp1* and *Atf4*, and the peak of *Krt8* expression (Fig. 6g). We additionally found a very high correlation of *Krt8* expression dynamics along the differentiation trajectory with the EGFR ligand amphiregulin (*Areg*), the Nuclear factor NF-kappa-B p100 subunit (*Nfkb2*), and the Tweak receptor CD266 (*Tnfrsf12a*), which is a TNF receptor superfamily (TNFSF) member possibly mediating non-canonical NF-kB signaling^54^ (Fig. 6g).

Finally, the transdifferentiation trajectory reached a critical pre-AT1 stage that was marked by the downregulation of the *Krt8* signature and the induction of a gene expression program reminiscent of the epithelial to mesenchymal transition (EMT), together with one of its master regulators *Sox4^55^* (Fig. 6g). We also observed pre-AT1 specific expression of the transcriptional coactivator TAZ (*Wwtr1*), which acts as a downstream regulatory target in the Hippo signaling pathway and was recently reported to be critical for AT2 cell differentiation during alveolar regeneration^17^. Also, the expression of the key downstream component of the canonical Wnt signaling pathway beta-catenin (*Ctnnb1*) transiently peaked in the pre-AT1 stage. Interestingly, we found many TGF-beta target genes, including *Sox4*, integrin beta-6 (*Itgb6*), and connective tissue growth factor (*Ctgf*) peaking in the pre-AT1 phase (Fig. 6g), suggesting that availability of TGF-beta is important for alveolar regeneration^18^. The non-muscle myosin heavy chain IIa (*Myh9*) also peaked in pre-AT1 cells, suggesting important additional cytoskeletal rearrangements and increased cell contractility in the already squamous Krt8+ alveolar progenitor cells in the final steps of maturation towards AT1 cells (Fig. 6g). Terminally differentiated AT1 cells were characterized by high expression of the transcription factors *Hopx, Gata6* and *Wwc1,* as well as a large number of developmentally important factors, including extracellular matrix proteins and growth factors, such as *Fgf1*, *Npnt* and *Agrn* (Fig. 6g).

We hypothesize that the Krt8+ alveolar progenitor cell state with its unique gene expression program serves important niche functions to coordinate other cell types during tissue regeneration. Thus, we leveraged the receptor-ligand database from the whole lung single cell survey (Fig. 1) to determine cell communication paths specific for Krt8+ progenitors in comparison to AT1. The largest number of receptor-ligand pairs was found with fibroblasts, macrophages and (capillary) endothelial cells (Fig. 6h; see Table S6 for all receptor-ligand pairs). AT1 cells are in tight contact with capillary endothelial cells, which enables their prime function of mediating gas exchange between inhaled air and blood. Thus, the connectome (all receptor-ligand interactions) of AT1 with endothelial cells is crucial for normal homeostasis. Interestingly, in the endothelial cell (EC) connectome with Krt8+ progenitors and AT1, the capillary endothelial cells received signals via the endothelin-receptor (*Ednrb*) expressed on EC via the ligand endothelin-1 (*Edn1*), which was specifically expressed on Krt8+ alveolar progenitors and not on AT1 (Fig. 6i.). Conversely, the AT1 cells displayed a large number of ligands, including *Vegfa* and *Sema3e* that bind receptors, such as *Flt1* or *Nrp1/2* on ECs, which were not expressed on Krt8+ progenitor cells. Similar selective differences between Krt8+ progenitors and AT1 were observed for receptors such as the urokinase plasminogen activator receptor (*Plaur*) specifically expressed on Krt8+ progenitors but not AT1, binding to the EC-derived ligand PAI-1 (Serpine1) (Fig. 6j).

Thus, we conclude that Krt8+ alveolar progenitor cells give rise to AT1 via a series of distinct transcriptional programs in a temporal order that we resolve in our analysis (Fig. 7). Krt8+ alveolar progenitors feature highly distinct cell-communication pathways compared to AT1. Based on the fact that the Krt8+ cell state precedes the formation of AT1 and its temporal co-occurrence with processes such as angiogenesis, fibrogenesis, and inflammation during the first three weeks of tissue repair, we propose critical importance of Krt8+ alveolar progenitors as instructive niche cells for these processes.

**Figure 7.**
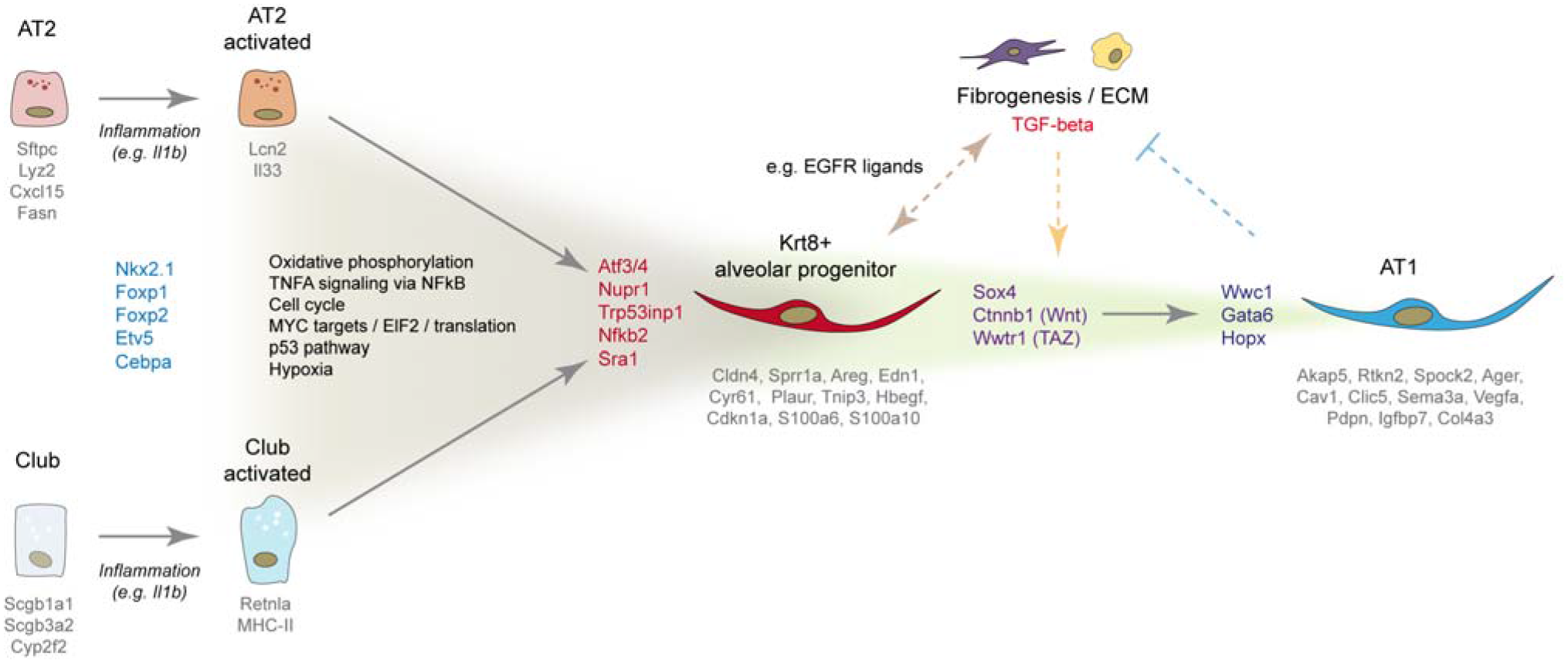
Summary of observations and revised model of alveolar regeneration. AT2 and club cells enter an activated state characterized by a distinct set of pro-inflammatory genes as a result of inflammatory cytokine driven NFkB activation early after injury. Both club and AT2 cells can converge on the same Krt8+ alveolar progenitor cell state as a consequence of metabolic reprogramming and the local niche in injured alveoli. Activated club and AT2 cells lose cell identity genes such as *Cebpa* and *Etv5* and re-enter the cell cycle, followed by a Myc-driven phase of cell growth and differentiation that is marked by increased oxidative phosphorylation and a drastic change in shape towards a squamous morphology. The Krt8+ alveolar progenitors are characterized by a stress response gene expression program involving p53, hypoxia response, and ER-stress related pathways. Terminal differentiation into AT1 cells involves a transient peak of the transcriptional regulators *Sox4*, *Ctnnb1* and *Wwtr1*, and their target genes. This indicates involvement of the Wnt, Yap/Taz, and TGF-beta signaling pathways in committing Krt8+ alveolar progenitors towards AT1 cell fate, which then becomes fixed by a peak in *Hopx* expression. Krt8+ alveolar progenitors peak in numbers during fibrogenesis and feature a highly distinct connectome of receptor-ligand pairs with endothelial cells, fibroblasts, and macrophages, suggesting that these cells serve to instruct the fibrogenic phase of lung regeneration. Conversely, it is likely that the fibrogenic niche, which as we show is largely driven by fibroblasts and macrophages, plays an instructive role in terminal AT1 differentiation. Finally, mature AT1 cells likely provide signals important for resolution of fibrogenesis. We generated a comprehensive list of regulated genes and receptor-ligand pairs resolved by time and cell type/state that can be accessed in a user-friendly interactive webtool.

## Discussion

In this work, we describe the dynamics of mouse lung regeneration at single cell resolution and discover the alveolar progenitor cell state that gives rise to AT1 cells. This unique cell state was marked by Keratin-8 expression and appeared transiently with its peak during the fibrogenic phase. We used a staged approach by first conducting a ‘bird’s-eye’ experiment surveying whole-lung single cell suspensions at 6 timepoints, followed by a targeted ‘sky dive’ analysis^31^ of epithelial cells with a higher longitudinal resolution at 18 timepoints after injury. Leveraging the power of pseudotemporal modeling^20,21,56,57^, we analyzed gene regulation during epithelial transdifferentiation, laying out the sequence of gene expression programs and highlighting key transcriptional regulators. Our inferred cell fate model was validated by correspondence with the real timepoints of sampling, RNA velocities of individual cells and lineage tracing experiments. The receptor-ligand analysis revealed potential routes of cell-cell communication and their dynamics over time. All data and code is freely available at our interactive webtool and github repository.

Recent evidence shows a high degree of plasticity between epithelial cell types^5^, including the possibility that the adult regenerative response may induce unique cell fate trajectories that do not resume developmental trajectories. Remarkably, even injury-induced reprogramming of differentiated cell types to bona fide long-lived stem cells has been demonstrated in the lung^58^ and other organs^59^. The new tools of scRNA-seq and computational lineage reconstruction employed here enable an unbiased assessment of cell plasticity after injury. Our analyses provide evidence that transdifferentiation processes are frequent and highly non-gradual, entailing intermediate gene expression programs that are distinct from both the initial and terminal cell states. In the classical model of stem cell differentiation, cells gradually lose stem cell identity genes and gain genes of the terminal state. In the Waddington landscape model^23^, such gradual differentiation of cells is described as a ball rolling down the hill (stem cell state) towards a valley (differentiated cell state). By contrast, in our transdifferentiation model presented here, the Krt8+ alveolar progenitor cell state represents a semi-stable plateau on a hill that cells need to ‘climb up’ to get from one cell state ‘valley’ (e.g. AT2) to the next cell state ‘valley’ (e.g. AT1).

Various potential stem and progenitor sources for AT1 cells have been described, including AT2 cells, bronchioalveolar stem cells (BASC) and p63(+)Krt5(+) distal airway stem cells (DASC). We did not see an important contribution of Krt5+ cells in our data, however, we observed a substantial fraction of airway-derived Krt8+ cells (Sox2-Cre). Our data revealed that activated club cells, in particular a previously unrecognized MHC-II subset, and activated AT2 cells gave rise to Krt8+ alveolar progenitor cells. Because some of the club cells co-expressed *Sftpc*, a contribution of BASCs cannot be excluded. It is remarkable that both AT2 cells and airway-derived progenitors within the club cell lineage converged on the same Krt8+ transcriptional state for their transdifferentiation towards AT1. This intriguing finding suggests that not the cellular lineage, but the injured alveolar niche determines (progenitor) cell identity. Along these lines, a recent scRNA-seq study of *C. elegans* embryogenesis showed that while in the early embryo the lineage-transcriptome correlation was high, in the final stages of development the distinct developmental lineages converged on identical transcriptional states if they migrated to the same location in the adult worm^60^. Similarly, we speculate that airway cells migrating towards injured alveoli and activated AT2 cells converge on the Krt8+ progenitor state because of the highly specific environment in the injured alveolar niche.

Many pathways that peaked in the Krt8+ progenitor cell state represent environmental stress- and inflammation-induced gene expression programs, represented by transcriptional master regulators, such as *Trp53*, *Atf3/4*, *Nupr1, Hif1a* and *NFkB*. This indicates that inflammation and changes in the cellular environment, including the hypoxic conditions present after lung injury^14^, may at least partially drive the generation of Krt8+ alveolar progenitors. Indeed, the proliferation of AT2 cells after lung injury was lost in AT2-specific IL1-receptor knock-out mice and involves Il1-beta and Tnf-alpha driven NFkB activation^16^, which provides a molecular link between inflammation and epithelial regeneration that is consistent with our results. We propose that environmental stress, together with inflammatory stimuli can promote cell plasticity by inducing epithelial cell states with a higher susceptibility for alternative fate programs. It has long been noted that isolated AT2 cells spontaneously drift towards AT1 fate *in vitro*, suggesting that plasticity may be a cell-intrinsic property and that AT2 cell identity *in vivo* is actively maintained by niche signals. Interestingly, during a five-day AT2 to AT1 *in vitro* differentiation, Krt8 protein levels were shown to be highest at day 3, followed by the AT1 marker Pdpn peaking later at day 5^61^.

In lung development, the generation of the distal epithelium has been proposed to be driven by a bipotent progenitor co-expressing both AT1 and AT2 markers^62^. Additionally, a recent scRNA-seq analysis of the mouse lung epithelium at birth identified a similar AT1/AT2 cluster that may be interpreted as a bipotent progenitor state^63^. In our analysis, both published developmental signatures do not correspond well to the injury induced Krt8+ alveolar progenitor signature described here. Interestingly, we do however find rare Krt8 high alveolar cells in the parenchyma of the normal adult mouse lung. Additional experiments will be needed to better understand the differences of epithelial lineage trajectories in lung development versus adult homeostasis and regeneration. Since Krt8+ alveolar progenitor cells do not express high levels of club, AT2 or AT1 cell marker genes, we consider them as a distinct ‘reprogrammed’ cell type entity with a high progenitor potential for AT1 cells. We cannot fully exclude that the Krt8+ progenitors also give rise to AT2 cells, however, our analysis suggests a strong bias towards AT1 cell differentiation.

Morphologically, the terminal differentiation of AT1 cells in development has been shown to occur via a non-proliferative two-step process of cell flattening and cell folding^9^. We have shown that Krt8+ alveolar progenitor cells feature mostly squamous morphology and may thus correspond with this first phase of cell flattening. In the developmental cell folding phase, AT1 cells increase their size ten-fold to span multiple alveoli and establish the honeycomb alveolar structure in coordination with myofibroblasts and capillary vessels^9^. In this process, AT1 cells express a large number of morphogens, such as *Vegfa* and semaphorins that stimulate angiogenesis and thus likely play an active signaling role in the coordination of alveolar morphogenesis. We confirm the specific expression of these morphogens only in mature AT1 in our study and show in contrast that Krt8+ progenitors express a distinct set of morphogens, including Endothelin-1 (*Edn1*) for the paracrine stimulation of capillary endothelial cells.

Our results predict distinct transcriptional regulators as key switch points in terminal AT1 differentiation, including TAZ (*Wwtr1*), *Sox4* and beta-catenin (*Ctnnb1*). The mechanistic importance of *Wwtr1* (YAP/TAZ - Hippo pathway) in AT2 to AT1 transdifferentiation has recently been demonstrated by using small molecule inhibition and conditional knock-out in mouse lung organoids and in vivo injury experiments^17,64^, thus validating our prediction that this factor commits Krt8+ progenitors for AT1 cell fate. Moreover, an important role of beta-catenin and the canonical Wnt signaling pathways has been suggested based on *in vitro* differentiation of isolated AT2 cells^61^. A functional role of *Sox4* in switching towards AT1 fate as suggested by our model is novel and awaits experimental validation. Furthermore, the hierarchy and crosstalk of the TGF-beta, Wnt and Hippo pathways in the final steps of Krt8+ progenitor commitment towards AT1 is still unclear and requires further dissection.

This work shows that injury induced epithelial cell plasticity in the lung involves a non-gradual transdifferentiation, leading to an intermediate and highly distinct progenitor cell state that requires a defined microenvironment to commit towards AT1 cell fate. We propose that the molecular definition of such intermediate progenitor cell states, as exemplified in our study, is highly relevant for understanding human diseases and is difficult to observe using bulk analysis methods. It is possible that in pathological scenarios, including chronic lung disease, such intermediate progenitor cell states may persist if signals for terminal differentiation are perturbed. Indeed, we found high amounts of KRT8+ cells in the parenchyma of interstitial lung disease patients. However, it is currently unclear whether their transcriptomic signature exactly compares to the murine Krt8+ alveolar progenitor cells described here. Co-expression of AT1 and AT2 markers in abnormal epithelial cells in idiopathic pulmonary fibrosis patients, together with aberrant activity of p53, TGF-beta, Hippo and Wnt pathway genes has been reported^65^. Furthermore, local hypoxia signaling has been implicated in dysplastic abnormal epithelial barriers^14^, which we suggest may represent an accumulation of intermediate progenitor cell state blocked in its commitment towards AT1 cell fate.

Ongoing efforts to chart the Human Lung Cell Atlas aim at improving the understanding of human lung development, adult homeostasis and regeneration as a basis to explore chronic lung disease^1^. A major difficulty in analyzing such dynamic processes in human tissues is the lack of longitudinal data. Thus, we believe that efforts in data integration across a large number of individuals^66^, together with perturbation experiments in human *ex vivo* tissue culture and organoid models will be needed to reach this aim. As shown in our work, mouse models of lung injury and disease can provide longitudinal data with high temporal and cellular resolution. Together with novel computational modeling approaches^67^, these data may enable mapping of dynamic processes across species, individuals, and experiments, thereby empowering future mechanistic predictions for human disease.

## Extended data Figures

**Figure S1.**
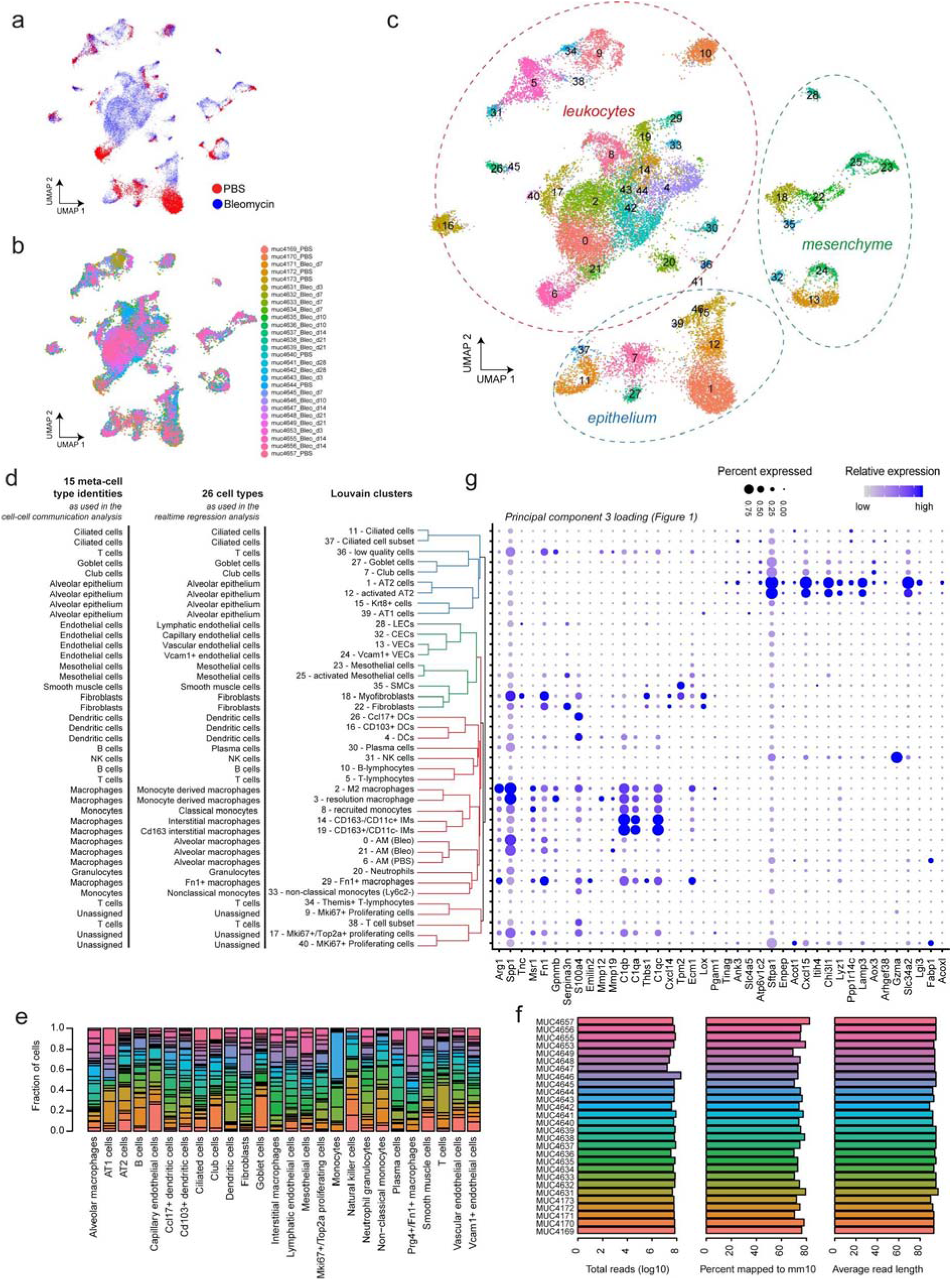
Good technical agreement of whole lung single cell transcriptomes of 28 individual mice. UMAP embeddings show good overlap between treatment conditions (a) and individual mouse replicates (b). (c) UMAP embedding colored by Louvain clusters demonstrates separation of cells into major lineages. (d) Unsupervised hierarchical clustering of the Louvain clusters recapitulates known hierarchical cell type topology. (e) Bar plot shows high overlap of mouse samples across cell types. (f) Alignment summary statistics are comparable across mouse samples. (g) Dot plot shows average expression of genes with top PC3 loadings across cell type identities.

**Figure S2.**
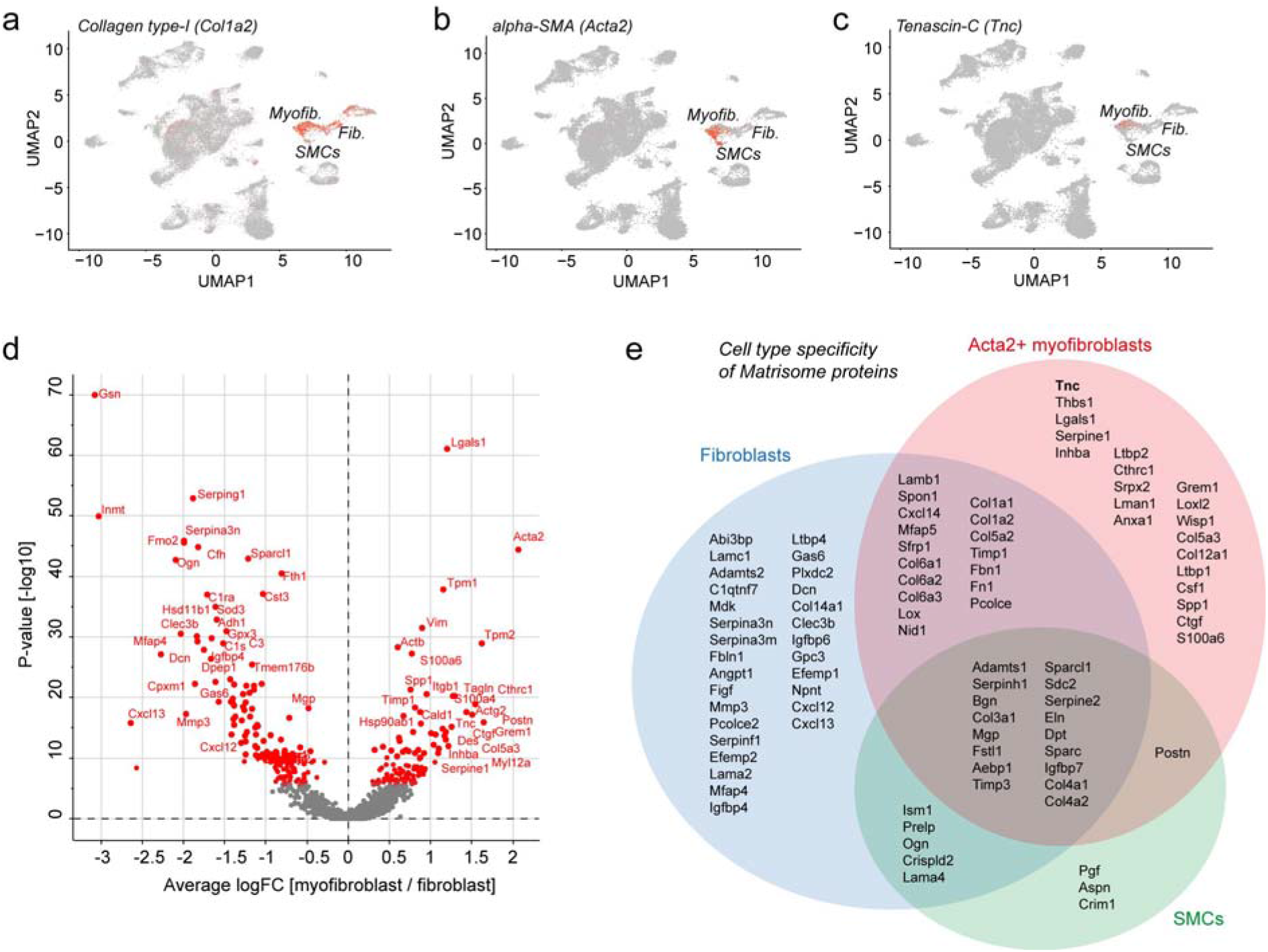
Transient appearance of the myofibroblast cell state upon lung injury. (a-c) Relative expression levels of *Col1a2* (a), *Acta2* (b), and *Tnc* (c) are shown on the UMAP embedding. (d) The volcano plot shows differential gene expression between myofibroblasts (right side) and fibroblasts (left side). (e) Single cell analysis was used to derive the myofibroblast specific ECM components in comparison to fibroblasts and smooth muscle cells.

**Figure S3.**
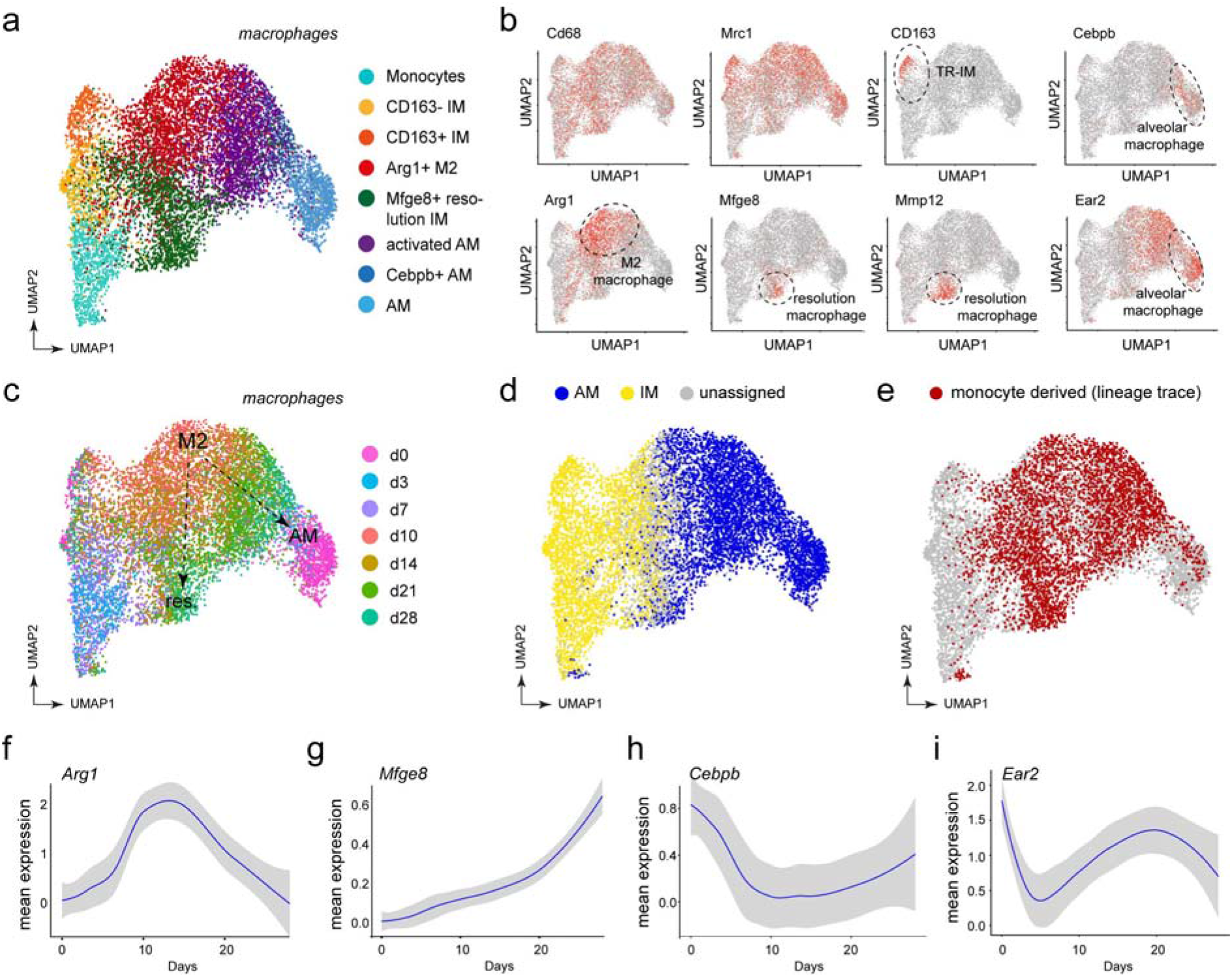
Dynamics of macrophage states in lung tissue regeneration. (a, c) UMAP embedding of 10379 cells that express known macrophage markers is colored by (a) cluster identity and (c) time points. Following cells along the time course after reaching the peak of inflammation at day 10 and 14, two potential trajectories can be discerned. (b) Several macrophage populations can be identified. These clusters uniformly express the macrophage marker *Cd68* and *Mrc1* while also showing distinct expression of certain genes. (d) Previously published gene signatures from bulk RNA experiments were used to reveal potential origins of macrophage cells. In this data set, FACS-sorting allowed to differentiate between tissue-resident alveolar (AM), interstitial (IM) and monocyte-derived macrophage populations^37^. Similarity score of each cell is calculated as correlation to differentially expressed genes and corresponding log fold changes in the three sorted populations. Cells are assigned to either AM or IM category, if the difference in scores for either category is higher than 0.05. Alveolar macrophages in our data set indeed show the highest score on the tissue-resident AM. (e) Potentially monocyte-derived cells based on scoring (at threshold of 0.1). There is a separation in the potentially monocyte-derived cells, which concurs with the real-time trajectories in (c) Relative cell type frequency per time point of (f) M2 macrophages, (g) Mfge8+ macrophages, and (h, i) alveolar macrophages across all samples and the smoothed expression per time point of (h) *Arg1*, (i) *Mfge8*, (h) *Cebpb*, and (i) *Ear2* in the macrophage subset with confidence interval of 0.95.

**Figure S4.**
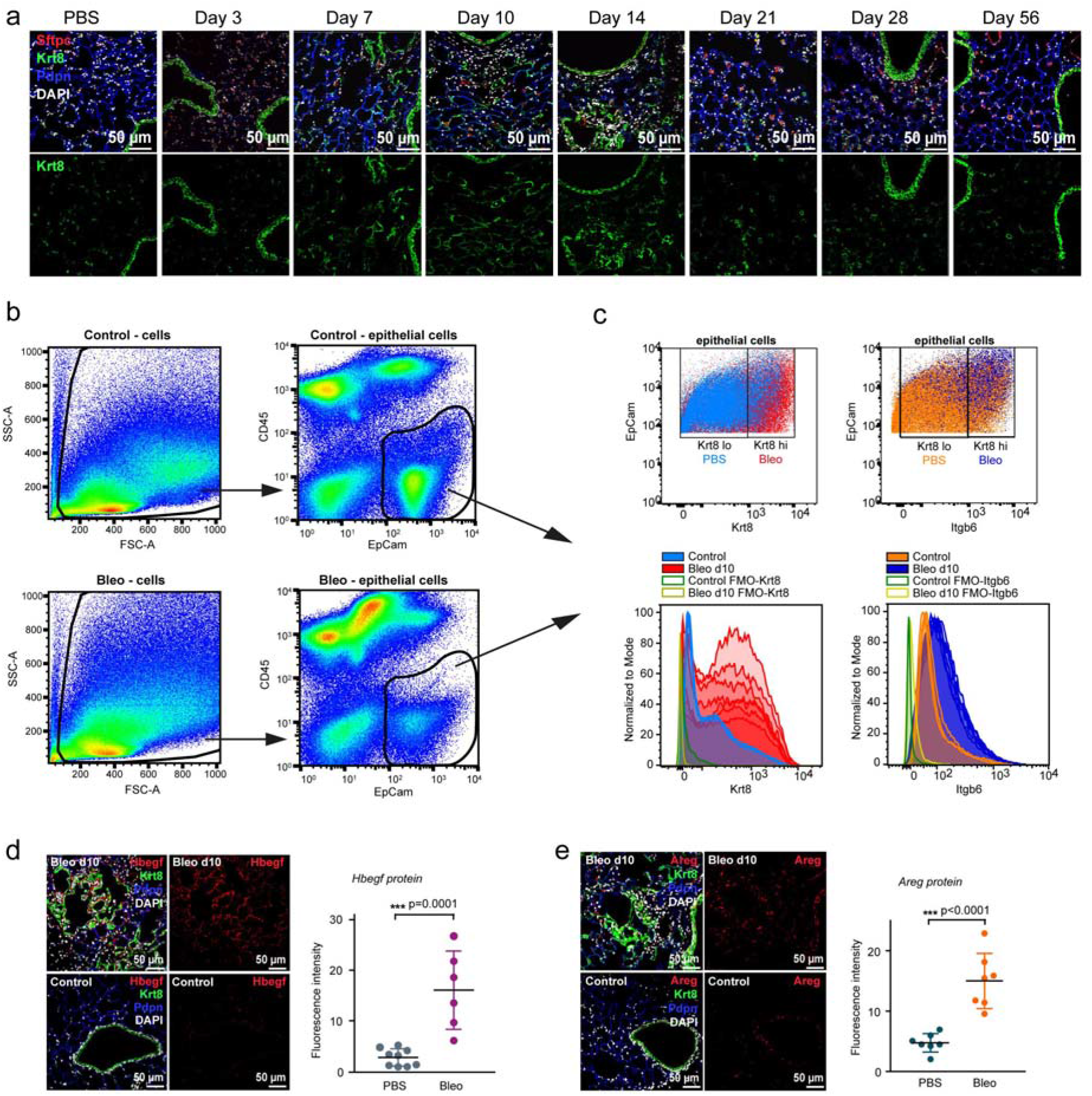
Protein validation of the alveolar Krt8+ cell signature. (a) Immunostaining of Krt8 (green) at the indicated time points after bleomycin injury. FFPE tissue sections were co-stained with the AT2 marker Sftpc (red), and the AT1 marker Pdpn (blue). Nuclei were labeled using DAPI (white). Scale bar = 50 microns. (b) Gating strategy for the analysis of CD45-/Epcam+ epithelial cells. (c) The scatter plots and histograms show increased expression of Krt8 and Itgb6 at day 10 after bleomycin in Epcam+ epithelial cells. Highest expression of Itgb6 was observed on Krt8 high cells. Fluorescence-minus-one (FMO) controls were used for both the Krt8 and Itgb6 quantification. (d) Increased Hbegf (red) expression in bleomycin treated lung tissue, showing partial overlap with Krt8 (green) signal. Quantification of the mean fluorescence signal intensities confirmed increased Hbegf expression (unpaired t-test *** p = 0.0001). Sections were co-stained with Pdpn (blue); scale bar = 50 microns. (e) Immunostainings of Areg (red) and Krt8 (green) expression in the lung, co-stained with Pdpn (blue) and quantified by mean fluorescence intensity. Unpaired t-test *** p < 0.0001. Scale bar = 50 microns.

**Figure S5.**
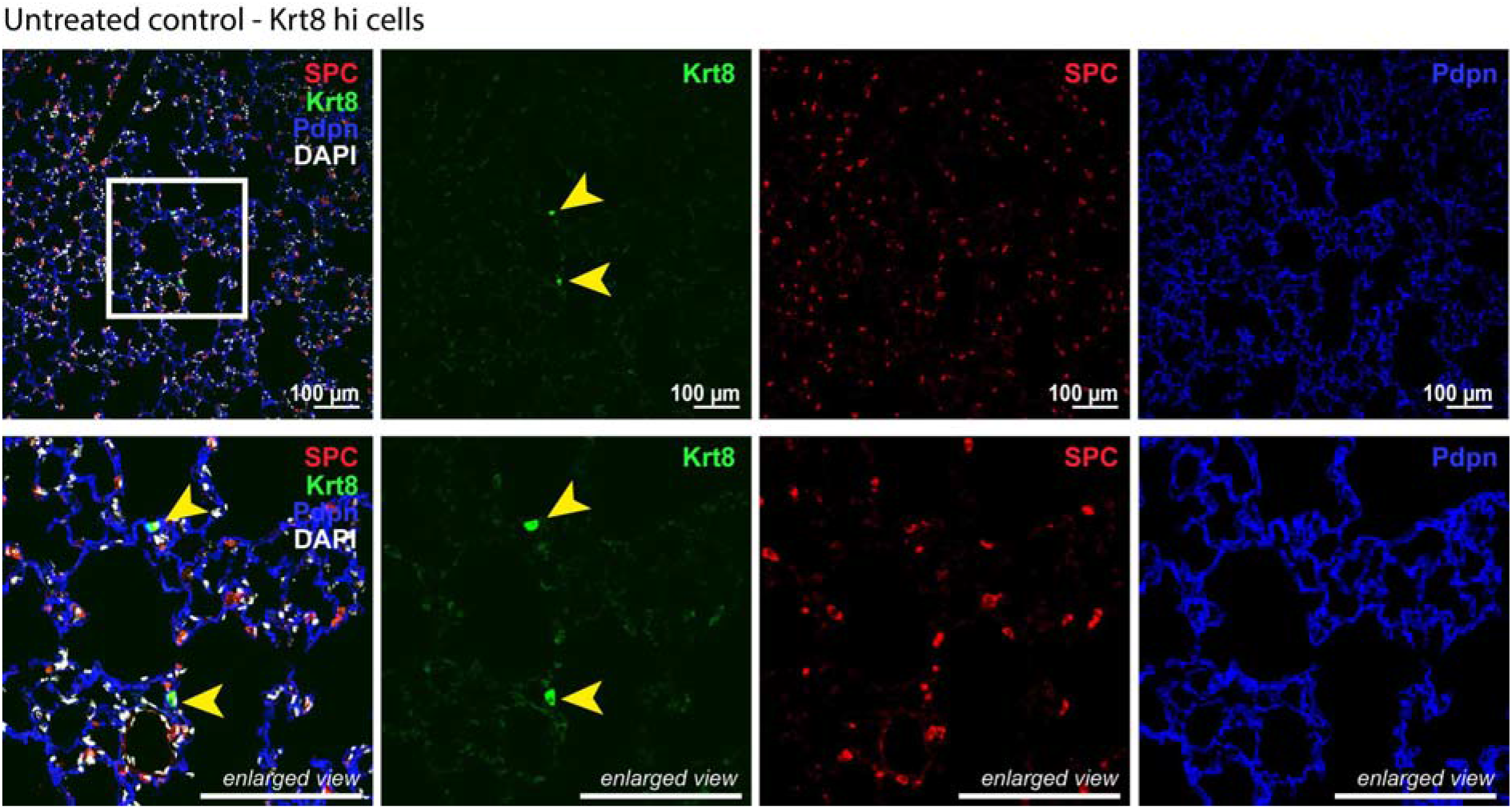
Rare Krt8+ cells in normal alveolar homeostasis. Fluorescent immunostainings and confocal imaging of lung sections from untreated control lungs. Nuclei (DAPI) are colored in white, Krt8 appears in green, Sftpc (AT2 cells) in red, and Pdpn (AT1 cells) in blue. The scale bar indicates 100 microns.

**Fig. S6.**
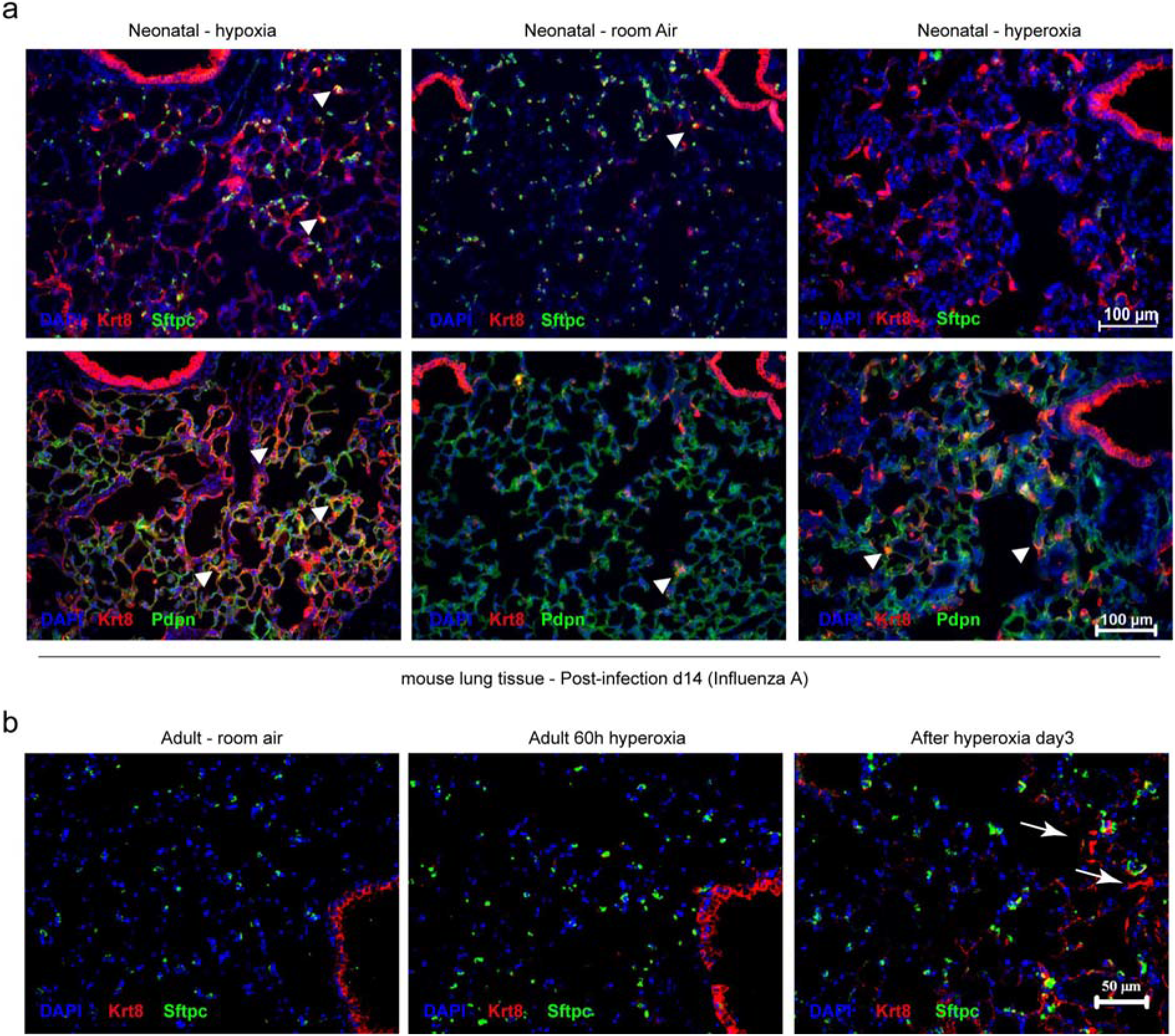
Appearance of Krt8+ alveolar progenitor cells in two alternative mouse injury models. (a) An aberrant oxygen environment at birth alters alveolar injury and repair following influenza A virus infection. Lungs of infected mice were stained for Krt8 (red) and Sftpc (green). Scale bar = 100 microns. (b) A sixty-hour exposure of adult mice to hyperoxia leads to the emergence of Krt8+ cells in the alveolar space. Mice were sacrificed three days after the exposure period terminated. Lung tissue was stained for Krt8 (red) and Sftpc (green). Scale bar = 50 microns.

**Figure S7.**
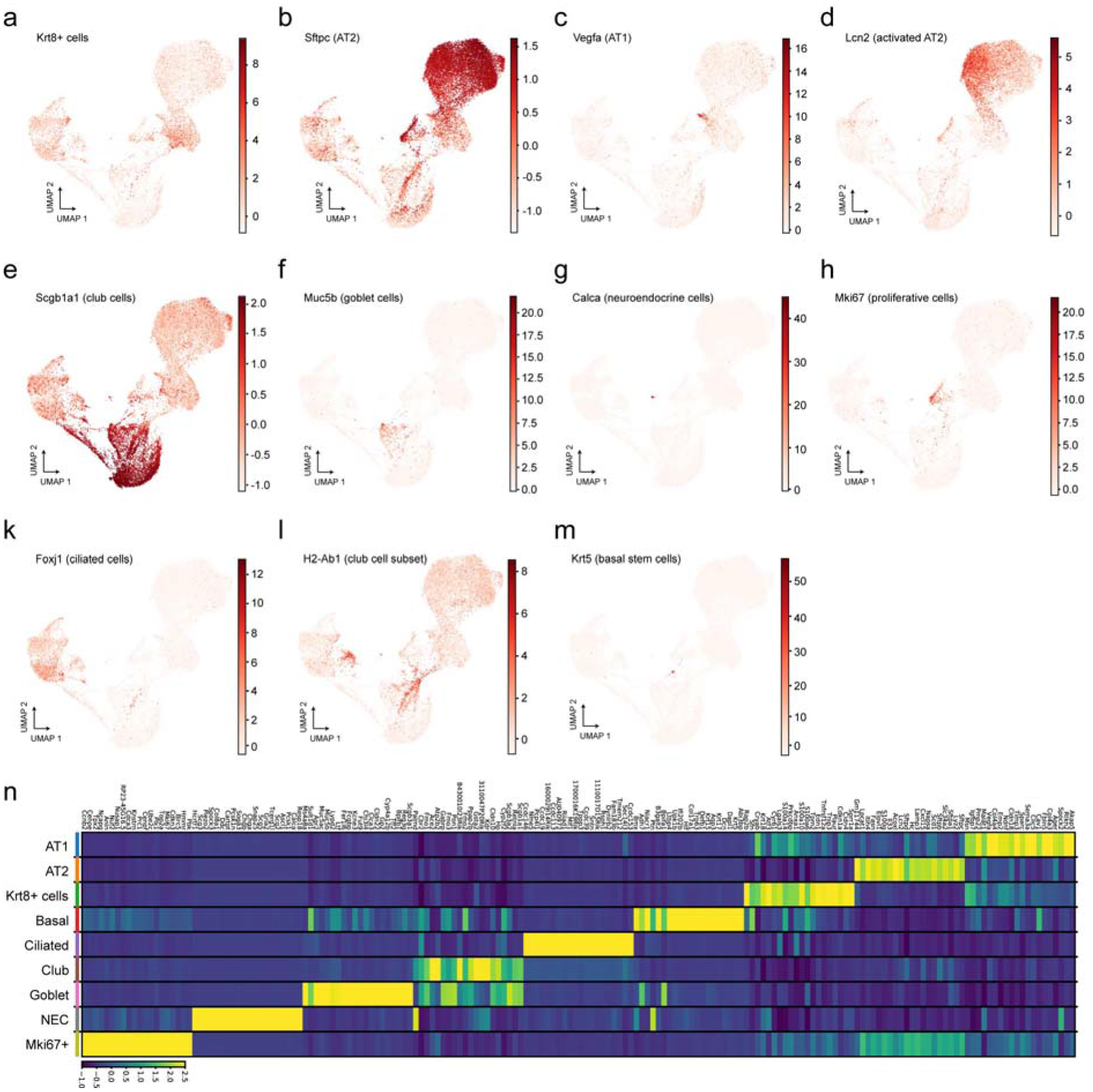
Feature plots of selected marker genes for epithelial cell types. UMAP embeddings display distinct expression patterns for selected epithelial cell type marker genes: (a) *Krt8* (ADI), (b) *Sftpc* (AT2 cells), (c)*Vegfa* (AT1 cells), (d) *Lcn2* (activated AT2 cells), (e) *Scgb1a1* (club cells), (f) *Muc5b* (goblet cells), (g) *Calca* (neuroendocrine cells), (h) *Mki67* (proliferative cells), (k) *Foxj1* (ciliated cells), (l) *H2-Ab1* (club cell subset), (m) *Krt5* (basal stem cells). Red colors indicate higher expression levels. (n) Heatmap shows the average expression levels for the top 20 genes with lowest adjusted p value of each meta cell type.

**Figure S8.**
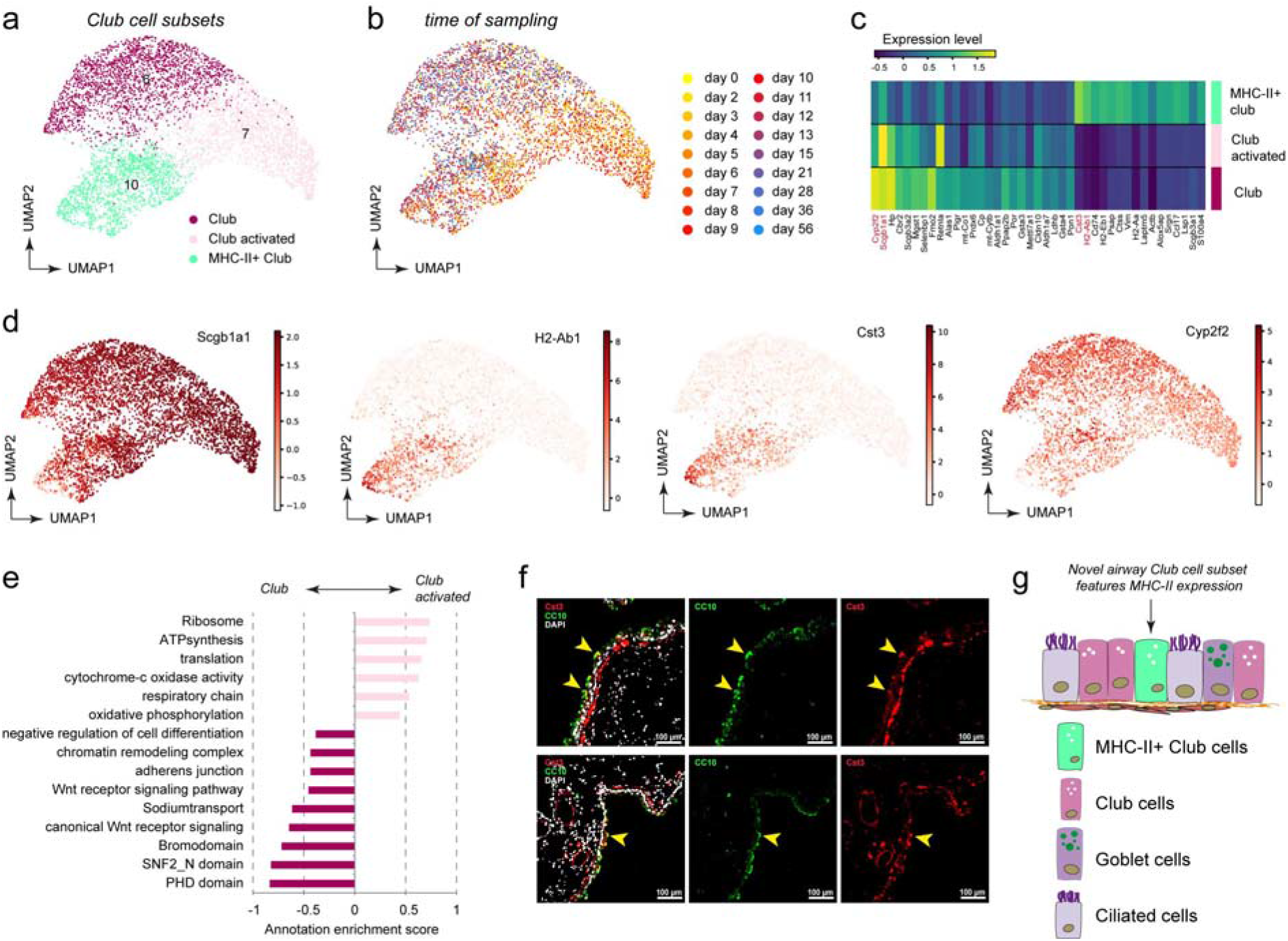
Discovery of a MHC-II positive club cell subset. Plots visualize the UMAP embedding of Club cells colored by Louvain clustering (a) and by time point (b). (c) The heatmap shows the average expression levels of marker genes across the three Club cell clusters. (d) UMAP embedding shows distinct expression patterns for selected marker genes. (e) The bar graph shows the annotation enrichment score^68^ for selected examples of gene categories with significant enrichment (FDR < 5%) in either activated Club (positive scores) or Club cells (negative scores). (f) Immunofluorescence staining of mouse airways shows CC10+ club cells (green) and Cst3+ cells (red), DAPI (white). Note the partial overlap of Cst3+/CC10+ airway cells (highlighted by yellow arrowheads). Scale bar = 100 microns. (g) Revised model of club cell heterogeneity in mouse airways.

**Figure S9.**
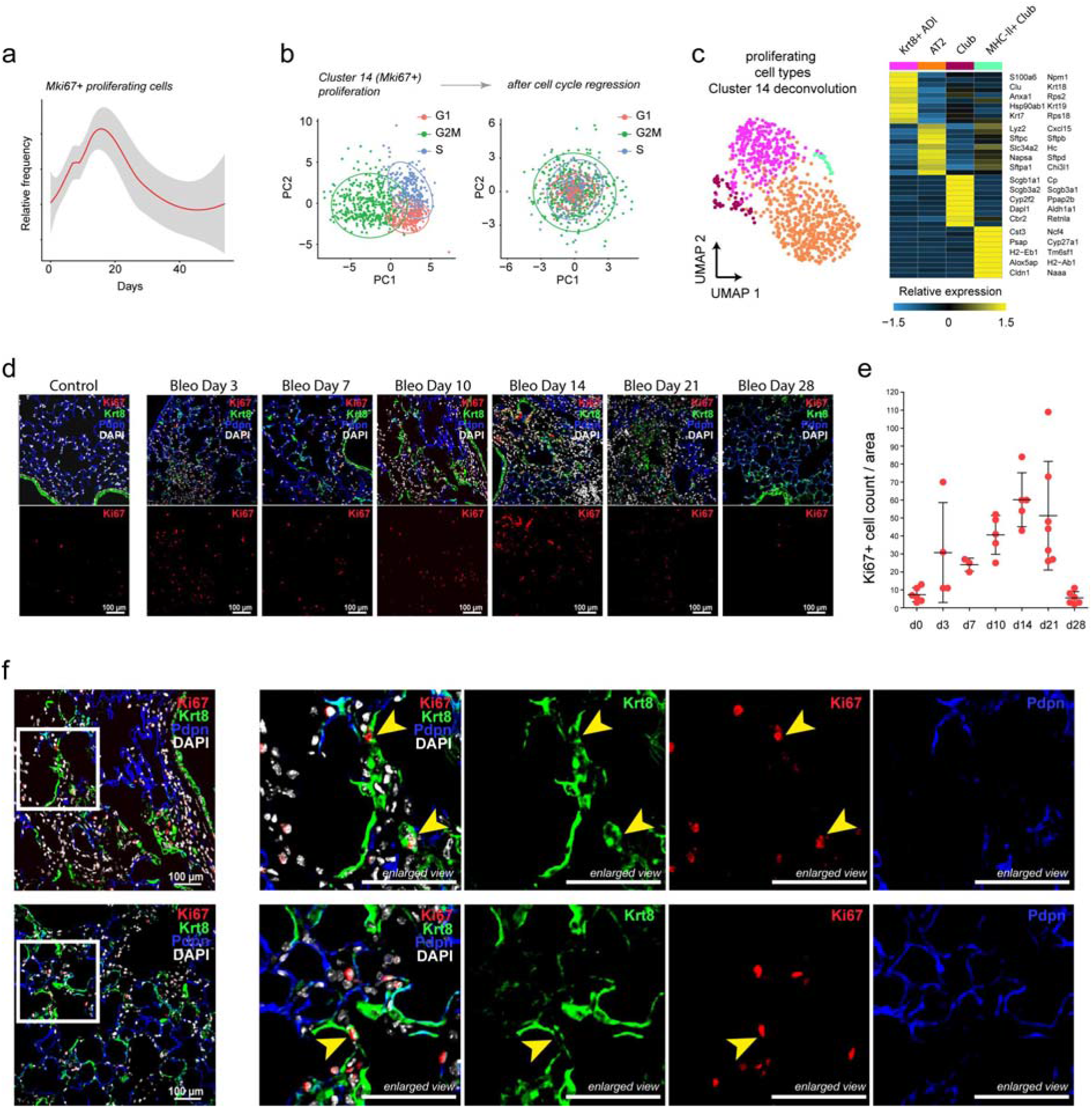
Cell cycle analysis reveals proliferation of Krt8+ alveolar progenitor cells. (a) Relative frequency of Mki67+ proliferating cells plotted over time. (b) The scatter plots show cells from proliferating cell cluster 14 before and after cell cycle regression, colored by inferred cell cycle phase. Regression removes cell cycle effects from principal component data manifold. Re-analysis of cell cycle corrected expression deconvolves cell type identities of proliferating cells. (c) UMAP of cell cycle corrected cluster 14 cells visualizes four distinct clusters, which contain Krt8+ progenitors, AT2, club, and MHC-II+ club cells. Heatmap shows the average expression levels of selected marker genes. (d) Immunofluorescence stainings of control versus bleomycin treated lung sections (day 3, 7, 10, 14, 21, 28). Sections were stained for Krt8 (green), Ki67 (red), Pdpn (blue), and DAPI (white). Scale bar indicates 100 microns. (e) Ki67+ cells were quantified from the micrographs by counting Ki67+ cells in each ROI/field of view (n=4-7 per time point, mean with SD). (f) Immunostaining as in (e) on day 10 post bleomycin injured lungs with enlarged views on proliferative Krt8+ alveolar progenitor cells (Ki67+/Krt8+), highlighted with yellow arrowheads. Scale bar indicates 100 microns.

## Methods

### Mouse experiments - bleomycin treatment

Pathogen-free female C57BL/6J mice were purchased from Charles River Germany and maintained at the appropriate biosafety level at constant temperature and humidity with a 12 hour light cycle. Animals were allowed food and water ad libitum. Animal handling, bleomycin/PBS administration, and organ withdrawal were performed in accordance with the governmental and international guidelines and ethical oversight by the local government for the administrative region of Upper Bavaria (Germany), registered under 55.2-1-54-2532-130-2014 and ROB-55.2-2532.Vet_02-16-208.

### Experimental design and animal treatment

Mice were divided randomly into two groups: (A) saline-only (PBS), or (B) bleomycin (Bleo). Lung injury and pulmonary fibrosis were induced by single-dose administration of bleomycin hydrochloride (Sigma Aldrich, Germany), which was dissolved in sterile PBS and given at 2U/kg (oropharyngeal instillation) and 3U/kg (intratracheal instillation) bodyweight. The control group was treated with sterile PBS only. Mice were sacrificed at designated time points (days 1-14, 21, 28, 35, 56) after instillation. Treated animals were continuously under strict observation with respect to phenotypic changes, abnormal behavior and signs of body weight loss.

### Generation of single cell suspensions from whole mouse lung tissue

Lung single cell suspensions were generated as previously described^24^. Briefly, after euthanasia, lung tissue was perfused with sterile saline through the heart and the right lung was tied off at the main bronchus. The left lung lobe was subsequently filled with 4% paraformaldehyde for later histologic analysis. Right lung lobes were removed, minced (tissue pieces at approx. 1 mm^2^), and transferred for mild enzymatic digestion for 20-30 min at 37°C in an enzymatic mix containing dispase (50 caseinolytic U/ml), collagenase (2 mg/ml), elastase (1 mg/ml), and DNase (30 μg/ml). Single cells were harvested by straining the digested tissue suspension through a 40 micron mesh. After centrifugation at 300 x g for 5 minutes, single cells were taken up in 1 ml of PBS (supplemented with 10% fetal calf serum), counted and critically assessed for single cell separation and overall cell viability. For Dropseq, cells were aliquoted in PBS supplemented with 0.04% of bovine serum albumin at a final concentration of 100 cells/μl.

### Production of microfluidic devices for Dropseq

Microfluidic devices needed for scRNAseq using the Dropseq platform were fabricated by means of standard soft lithography. In brief, by using photolithography, a polydimethylsiloxane (PDMS) master mold for the Dropseq device design (CAD file available as a download from: http://mccarrolllab.org/dropseq/) was fabricated from a SU-8 photoresist (MicroChem, USA), and spin-coated on a 3” silicon wafer to generate 125 μm-thick uniform layers. Afterwards, the master mold was filled with a 10:1 mixture of base to curing agent of the PDMS kit Sylgard 184 (Dow Corning, USA) and left at 60°C in an oven for 4 hours to crosslink the PDMS. After crosslinking, the PDMS replica was cut and peeled off from the master mold, as well as all necessary inlets/outlets for tubing connection were made in it using a 1 mm puncher. Next, the replica was sealed with a 2” x 3” microscopic slide, after the treatment of both in O_2_ plasma. The assembled microfluidic device was treated with Aquapel (Pittsburgh Glass Works, USA) to make all inner surfaces evenly hydrophobic.

### Single cell RNA-sequencing using Dropseq

Dropseq experiments were performed according to the original protocols^27,24^. Using the microfluidic device, single cells (100/μl) were co-encapsulated in droplets with barcoded beads (120/μl, purchased from ChemGenes Corporation, Wilmington, MA) at rates of 4000 μl/hr. Droplet emulsions were collected for 10-20 min/each prior to droplet breakage by perfluorooctanol (Sigma-Aldrich). After breakage, beads were harvested and the hybridized mRNA transcripts reverse transcribed (Maxima RT, Thermo Fisher). Unused primers were removed by the addition of exonuclease I (New England Biolabs), following which, beads were washed, counted, and aliquoted for pre-amplification (2000 beads/reaction, equals ca. 100 cells/reaction) with 12 PCR cycles (Smart PCR primer: AAGCAGTGGTATCAACGCAGAGT (100 μM), 2x KAPA HiFi Hotstart Ready-mix (KAPA Biosystems), cycle conditions: 3 min 95°C, 4 cycles of 20s 98°C, 45s 65°C, 3 min 72°C, followed by 8 cycles of 20s 98°C, 20s 67°C, 3 min 72°C, then 5 min at 72°C)^27^. PCR products of each sample were pooled and purified twice by 0.6x clean-up beads (CleanNA), following the manufacturer’s instructions. Prior to tagmentation, complementary DNA (cDNA) samples were loaded on a DNA High Sensitivity Chip on the 2100 Bioanalyzer (Agilent) to ensure transcript integrity, purity, and amount. For each sample, 1 ng of pre-amplified cDNA from an estimated 1000 cells was tagmented by Nextera XT (Illumina) with a custom P5-primer (Integrated DNA Technologies). Single-cell libraries were sequenced in a 100 bp paired-end run on the Illumina HiSeq4000 using 0.2 nM denatured sample and 5% PhiX spike-in. For priming of read 1, 0.5 μM Read1CustSeqB (primer sequence: GCCTGTCCGCGGAAGCAGTGGTATCAACGCAGAGTAC) was used.

Quality metrics, including the number of unique molecular identifiers (UMI), genes detected per cell and reads aligned to the mouse genome were comparable across all mice (Fig. S1). Every timepoint was analyzed together with control mice that were instilled with phosphate-buffered saline (PBS). UMI-based counting of mRNA copies was used to determine differential gene expression between single cells. We used the six batches of PBS control mice to exclude dominant batch effects observing very good overlap across mouse samples (Silhouette coefficient: −0.08) (Fig. S1).

### Processing of the whole lung data set

For the whole lung data set, the Dropseq computational pipeline was used (version 2.0) as previously described^32^. Briefly, STAR (version 2.5.2a) was used for mapping^69^. Reads were aligned to the mm10 reference genome (provided by the Dropseq group, GSE63269). For barcode filtering, we excluded barcodes with less than 200 detected genes. As 1000 cells were expected per sample, the first 1200 cells were used before further filtering. A high proportion (> 10%) of transcript counts derived from mitochondria-encoded genes may indicate low cell quality, and we removed these unqualified cells from downstream analysis. Cells with a high number of UMI counts may represent doublets, thus only cells with less than 5000 UMIs were used in downstream analysis.

### Analysis of the whole lung data set

The computational analysis of the whole lung data set was largely performed using the R package Seurat^70^. Count matrices were merged using Seurat version 2.3. The merged expression matrix was normalized using the Seurat NormalizeData() function. To mitigate the effects of unwanted sources of cell-to-cell variation, we regressed out the number of UMI counts using the Seurat function ScaleData(). Highly variable genes were calculated per sample, selecting the top 7000 genes with a mean expression between 0.01 and 8. After excluding homologs of known cell-cycle marker genes^71^, a total of 18893 genes were subjected to independent component analysis. The first 50 independent components were used as input to the FindClusters() function with the ‘resolution’ parameter set to two and the RunUMAP() function with the “n_neighbors” parameter set to ten.

#### Multi-omic data integration

To confirm global expression changes observed at the single-cell level, we integrated previously published bulk RNAseq and proteomics data obtained from whole mouse lungs 14 days after bleomycin-induced injury and controls^34^. Multi-omic data integration was performed as described previously^28,34^. Briefly, in silico bulk samples were generated by summing all counts within a mouse sample. Both the in silico bulk and whole lung tissue bulk data were normalized using the voom() function of the limma R package^72^. Next, in silico bulk, whole lung tissue bulk, and proteomics data were merged on a set of genes present in all three data sets and quantile normalized. This merged and quantile normalized expression matrix was then subjected to principal component analysis (PCA).

#### Discovery of cell type identity marker genes

To identify cluster-specific marker genes, the Seurat FindAllMarkers() function was applied, restricted to genes detected in more than 10% of cells and with an average fold change difference of 0.25 or more. Based on these derived marker genes and manual curation we assigned all clusters to cell type and meta-cell type identities (Fig. S1d). Cell type frequencies were calculated by dividing the number of cells annotated to a specific cell type identity, by the total number of cells for each mouse sample. In droplet-based scRNAseq data, background mRNA contamination by the so-called “ambient RNAs” is frequently observed. These mRNAs are believed to stem from dying cells which release their content upon cell lysis. This contamination is distributed to many droplets and leads to a blurred expression signal that does not stem solely from the single cell in the droplet but also from the solution that contains it. We used the function inferNonExpressedGenes() from SoupX^73^ to identify a set of 80 ambient RNAs and accounted for these in the downstream analysis.

#### Time course differential expression analysis

To identify genes that show differential expression patterns across time within a given cell type we performed the following analysis. We used the R packages splines and lmtest for our modeling approach. First, we manually combined the Louvain clusters into 26 cell types to generate a more coarse grained cell type annotation for the time course differential expression analysis (Fig. S1d). Within each of these groups we modeled gene expression as a binomial response where the likelihood of detection of each gene within each mouse sample was the dependent variable. Therefore, the sample size of the model was the number of mouse samples (n = 28) and not the number of cells. To assess significance we performed a likelihood-ratio test between the following two models. For the first model, the independent variables contained an offset for the log-transformed average total UMI count and a natural splines fit of the time course variable with two degrees of freedom. The independent variables of the second model just contained the offset for the log-transformed average total UMI count. The dependent variable of both models was the number of cells with UMI count greater than zero out of all cells for a given cell type and mouse sample. To account for potential false positive signal derived from ambient RNA levels, we calculated cell type marker genes for the 26 cell type annotation using the Seurat FindAllMarkers() function. For all 80 candidate ambient RNAs, we consequently set all regression p-values to one in cell types where the gene was not simultaneously a marker gene with an adjusted p-value of less than 0.1 and a positive average log fold change.

### Cell-cell communication analysis

To identify cell-cell communication networks, we downloaded a list of annotated receptor-ligand pairs^74^. Next, we integrated this information with the cell type marker genes from Table S1. Cell-cell communication networks were generated in the following manner. An edge was created between two cell types if these two cell types shared a receptor-ligand pair between their as marker genes.

#### Macrophage analysis

It is not entirely understood whether monocyte-derived macrophages contribute to the development of lung fibrosis. To see if our data reflects published models of monocyte recruitment, we integrated bulk RNAseq data from FACS sorted macrophage populations after bleomycin-induced lung fibrosis^37,75^. This data set contained bulk RNAseq gene expression of tissue-resident alveolar macrophages (TR-AMs), monocyte-derived alveolar macrophages (Mo-AMs), interstitial macrophages (IM), and monocytes (Mono) for both day 14 and day 19 after bleomycin injury, including additional measurements for TR-AMs at day 0. To derive a gene expression signature from the bulk RNAseq data, we used the R package limma^76^. We followed the standard limma workflow^75^ to find genes which are differentially expressed between these four populations. Next, we subset our scRNAseq data set to only clusters expressing known macrophage markers and selected a new set of variable genes. Following this the PCA and UMAPs were recreated for this subset, using 20 PCs and 20 n_neighbors in Seurat’s functions. The macrophages from our data were scored according to their similarity to these bulk-derived signatures using Pearson correlation. For each of the four bulk-derived groups, the log fold changes of the 500 most differentially expressed genes were correlated with the scaled expression values of each macrophage cell in our scRNAseq data. To separate potential monocyte-derived macrophages from interstitial macrophages, we assigned each cell to the category with the higher correlation coefficient as long as the difference was greater than 0.05. Otherwise, the cell was labeled ‘unassigned’.

### Processing of the high-resolution epithelial data set

The high-resolution gene expression matrix was generated as specified for the whole lung data set with the following changes. To lessen the technical bias introduced by ambient RNA, we applied SoupX the pCut parameter set to 0.3 within each sample before merging the count matrices together. The merged expression table was then pre-processed as described in the *Processing of the whole lung data set* section with minor alterations. To account for the fact that a certain fraction of the counts was removed, the upper threshold for the number of total UMI counts per cell was set to 3000.

### Analysis of the high-resolution epithelial data set

The computational analysis of the whole lung data set was performed using a combination of the Seurat^70^ and Scanpy^77^ code. Cell-cycle effects, the percentage of mitochondrial reads, and the total number of UMI counts are often viewed as unwanted sources of variation and were therefore regressed out using the Seurat functions CellCycleScoring() and ScaleData(). Genes which had a variable expression in at least two samples (17038 genes) were used for the principal component analysis. The majority of the cells were airway and alveolar epithelial cells, although non-epithelial cells were also captured. To filter the data further, the cells were clustered and clusters expressing non-epithelial markers were excluded from the data set. The cleaned object was then converted to a .h5ad file for downstream analysis using the python package Scanpy. The aligned bam files were used as input for Velocyto^19^ to derive the counts of unspliced and spliced reads in loom format. Next, the sample-wise loom files were combined, normalized and log transformed using scvelos (https://github.com/theislab/scvelo) functions normalize_per_cell() and log1p(). After merging the loom information to the exported .h5ad file using scvelos merge() function the object was scaled and the neighbourhood graph constructed with Batch balanced KNN (BBKNN)^78^ to account for the different PCR cycles used in the experiment with neighbors_within_batch set to 15 and n_pcs to 40. Two dimensional visualization and clustering was carried out with the Scanpy functions tl.louvain() at resolution two and tl.umap(). The neuroendocrine cells (NEC) formed a distinct cluster in the UMAP, however, they were only assigned to a single cluster at higher resolutions. To separate them from basal cells we captured the NEC with dbscan using the UMAP coordinates and assigned them as cluster 21. After manual curation of the markers the remaining 20 cluster were combined, leading to thirteen final meta cell types.

#### Cell-cycle analysis

The proliferating cells (Louvain cluster 14, Fig. 4d) of the high-resolution data set were subjected to cell type deconvolution analysis. Cell cycle phases (“S.Score”, “G2M.Score”) were regressed out using the Seurat ScaleData() function. Next, PCA was calculated using all unique marker genes from Table S3 and the Seurat RunPCA() function. UMAP embedding and Louvain clusters were calculated using the first 20 principal components with the Seurat RunUMAP() and FindClusters() functions, respectively. Upon manual curation of the marker genes for the generated embedding, we identified four distinct clusters. Next, the frequency of proliferating cells was calculated by dividing the number of cells in cluster 14, by the number of total cells for each mouse sample.

#### PAGA analysis

To assess the global connectivity topology between the Louvain clusters we applied Partition-based graph abstraction (PAGA)^42^. We applied the tl.paga() function integrated in the Scanpy package to calculate connectivities and used the Louvain clusters as partitions. The weighted edges represent a statistical measure of connectivity between the partitions. Connections with a weight less than 0.3 were removed.

#### Velocity analyses

To infer future states of individual cells we made use of the spliced and unspliced information. We employed *scvelo* (https://github.com/theislab/scvelo). The previously normalized and log transformed data was the starting point to calculate first and second order moments for each cell across its nearest neighbors (scvelo.pp.moments(n_pcs = 40, n_neighbors = 15)). Next, the velocities were estimated and the velocity graph constructed using the scvelo.tl.velocity() with the mode set to ‘stochastic’ and scvelo.tl.velocity_graph() functions. Velocities were visualized on top of the previously calculated UMAP coordinates with the scvelo.tl.velocity_embedding() function. To compute the terminal state likelihood of a subset of cells, the function scvelo.tl.terminal_states() with default parameters was used.

#### Trajectory differential expression analysis

To identify genes showing significantly altered expression across the differentiation trajectory towards the Krt8+ cell state, the following approach was used. The high-resolution data set was restricted to cells from Louvain clusters 2, 10 and 11. The dbscan() function from the DBSCAN R package was used to identify outlier cells which were subsequently removed from further analysis. The R package slingshot was used to infer the pseudotemporal ordering across the trajectory of the first two diffusion components of all remaining cells. Next, the analysis was restricted to genes with more than ten total UMI counts in more than five mouse samples. For each gene, the following generalized additive model was fitted using the R package gam. Gene expression was used as the explanatory variable and defined as a binary outcome representing the detection of the gene (UMI count > 0). Log-transformed total UMI counts were included as a covariate in the model to account for differences in library size. A smooth loess fit of the pseudotemporal coordinate was included as the second independent variable. As the interpretation of p-values across pseudotime is difficult, we defined all genes with a marginal p-value of less than 1e-5 as significant. Gene expression patterns along pseudotemporal trajectories were visualized using local polynomial regression fitting as implemented in the R loess() function with default parameters.

### Pathway analysis

To predict the activity of pathways and cellular functions based on the observed gene expression changes, we used the Ingenuity Pathway Analysis platform (IPA, QIAGEN Redwood City, www.qiagen.com/ingenuity) as previously described^34^. The analysis uses a suite of algorithms and tools embedded in IPA for inferring and scoring regulator networks upstream of gene-expression data based on a large-scale causal network derived from the Ingenuity Knowledge Base.

### Magnetic-activated cell sorting (MACS)

Cells from whole lung single cell suspensions were strained using a 40 μm mesh size and red blood cells were eliminated by lysis (RBC lysis buffer, ThermoFisher). For positive epithelial cell selection, cells were stained with CD326-AlexaFluor647 antibody (Biolegend, 118212) for 30 min at 4°C in the dark, and after washing, incubated with microbeads specific against AlexaFluor647 (Miltenyi Biotec, 130-091-395) for 15 min at 4°C. MACS LS columns (Miltenyi Biotec, 130-042-401) were prepared according to the manufacturer’s instructions. Cells were applied to the columns and positively-labeled epithelial cells were retained in the column. The flow-through was collected separately for later mesenchymal cell enrichment (negative MACS selection) and kept on ice. Epithelial cells were eluted from the LS columns and used for either Dropseq runs. Mesenchymal cells from the flow-through were further enriched by negative depletion of CD31+ (Invitrogen, 17-0311-82), CD45+ (Biolegend, 103112), Lyve1+ (Invitrogen, 50-0443-82), Ter119+ (Biolegend, 116218), and CD326+ cells (Biolegend, 118212). After antibody staining, 100 μl per 10 million cells of MACS dead cell removal beads (Miltenyi Biotec, 130-090-101) were added and incubated according to the product’s accompanying protocols. Depletion of undesired cell types was achieved by the use of microbeads specific for APC (Miltenyi Biotec, 130-090-855), which ensured magnetic retention of these cells. Likewise to epithelial cells, negatively-selected mesenchymal cells were applied to the Dropseq workflow.

### Flow cytometry

Isolated total lung cell suspensions were used to detect and quantify cell populations by flow cytometry. After depletion of red blood cells by red blood cell lysis buffer (Invitrogen, ThermoFisher), cell suspensions were stained with anti-mouse CD45-PE-Vio770 (Miltenyi Biotec, 130-110-661), CD326-BV421 (Biolegend, 118225), Krt8/TROMA-I (DSHB-Developmental Studies Hybridoma Bank at the University of Iowa), and αvβ6-specific monoclonal antibody 6.3G9 (Itgb6-3G9; kindly provided by Prof. Dr. Dean Sheppard, available through Biogen Idec, USA). Cells were stained for surface markers in the dark at 4°C for 20 min, followed by cell fixation and permeabilization (Fix & Perm, Life Technologies, GAS004) for intracellular staining of Krt8. Epithelial cells were selected using the CD45-negative fraction of the cell isolate that stained positively for CD326. Within the epithelial cell gate, Krt8+, Itgb6+, or Krt8+/Itgb6+ cells were identified and quantified by their geometric mean fluorescence signal intensity. For exclusion of non-specific antibody binding and autofluorescence signal, fluorescence minus one (FMO) controls were included in the measurement. All stainings were performed per 1,000,000 cells in the following dilutions: CD326 (1:500), CD45 (1:20), Krt8 (1:35), Itgb6 (1:1000). Data was acquired in a BD LSRII flow cytometer (Becton Dickinson, Heidelberg, Germany) and analyzed by mean fluorescence intensity (MFI) using the FlowJo software (TreeStart Inc., Ashland, OR, USA). Negative thresholds for gating were set according to isotype-labeled and unstained controls.

### Precision cut lung slices (PCLS)

Precision cut lung slices were generated as previously described^79^. Briefly, using a syringe pump, the mouse lungs were filled via a tracheal cannula with 2% (w/v) warm, low gelling temperature melting point agarose (Sigma Aldrich, A9414) in sterile DMEM/Ham’s F12 cultivation medium (Gibco, 12634010), supplemented with 100 U/ml penicillin, 100 μg/ml streptomycin, and 2.5 μg/ml amphotericin B (Sigma Aldrich, A2942). Afterwards, the lungs were removed and transferred on ice in cultivation medium for 10 min to allow for gelling of the agarose. Each lung lobe was separated and cut with a vibratome (Hyrax V55; Zeiss, Jena, Germany) in 300 μm thick sections. The PCLS were immediately fixed in −20°C-cold methanol for 20 min and subsequently stained for immunofluorescence microscopy.

### Human lung material

For immunofluorescence stainings, resected human lung tissue and lung explant material was obtained from the CPC-M bioArchive at the Comprehensive Pneumology Center (CPC), Munich. ILD diagnosed lung tissue (n=5) is derived from lung explant material from transplantations, reflecting non-resolving end-stage disease. The tissue sections from patients with ARDS (n=2) have been provided by the Institute of Pathology at the University Hospital of Ludwigs Maximilians University, Munich. Healthy control tissue (n=7) was derived from tumor resections. All participants gave written informed consent. Tissue handling and this study were performed in accordance with the guidelines approved by the local ethics committee of the Ludwig Maximilians University, Munich, Germany (333-10).

### Immunofluorescence staining of PCLS, 3D microscopy and 3D morphometric analysis

Methanol-fixed PCLS were stained and imaged as previously described^80^. Shortly, primary antibodies were diluted in 1% bovine serum albumin (BSA, Sigma Aldrich, 84503) in PBS (1:100), incubated for 16 hours at 4°C and subsequently washed three times with PBS for 5 minutes each. Secondary antibodies were diluted in 1% bovine serum albumin in PBS (1:200), incubated for 4 hours at room temperature and subsequently washed three times with PBS for 5 minutes each. Primary antibodies were: rat anti-Krt8/TROMA-I (1:200; DSHB-Developmental Studies Hybridoma Bank at the University of Iowa), rabbit anti-pro-SPC (1:200; Millipore, AB3786), goat anti-Pdpn (1:200; R&D Systems, AF3244). Cell nuclei were stained with DAPI (40,6-diamidino-2-phenylindole, Sigma-Aldrich, 1:2,000). Confocal high-resolution 3D imaging of the PCLS was accomplished by placing the PCLS into a glass-bottomed 35 mm CellView cell culture dish (Greiner BioOne, 627870) as a wet chamber. Images of PCLS were acquired as z-stacks using an inverted microscope stand with an LSM 710 (Zeiss) confocal module operated in multitrack mode using the following objectives: Plan-Apochromat W 40x/1.0 M27 and Plan-Apochromat W 63x/1.3 M27. The automated microscopy system was driven by ZEN2009 (Zeiss) software. The acquired confocal fluorescent z-stacks were surface rendered in Imaris 9.3 software (Bitplane) and its statistical analysis tool (Measurement Pro) was used for 3D cell shape analysis using morphometric parameter sphericity as a readout (a value of 1 corresponds to a perfect sphere).

### Immunofluorescence microscopy

After euthanasia, mouse lungs were immediately inflated with 4% paraformaldehyde. For frozen OCT embedding, tissue was fixed for 1h at room temperature. Thin lung sections (7 μm) were cut on a cryostat. Sections were incubated with 0.1% sodium borohydride (PBS) to reduce background fluorescence, followed by blocking in PBS plus 1% bovine serum albumin, 5% non-immune horse serum (UCSF Cell Culture Facility), 0.1% Triton X-100 (Carl Roth, 3051.3) and 0.02% sodium azide (Sigma Aldrich, S2002). Slides were then incubated in primary antibodies overnight at 4°C followed by secondary antibody incubation at 1:1,000 dilutions at room temperature for >1 hr. Slides were counterstained with 1 μM DAPI for 5 minutes at room temperature and mounted using Prolong Gold (Life Technologies, P36930). The following antibodies were used: rabbit anti-pro-SPC (1:2,500; Millipore, AB3786), and rat anti-Krt8 (0.9μg/ml; TROMA-I (Krt8); University of Iowa Hybridoma Bank). Slides were imaged using a Leica Microscope (DM6B-Z; Leica Biosystems) or Axivision Imager M1 (Carl Zeiss AG).

For formalin-fixed, paraffin-embedded (FFPE) lung tissue, sections were cut at 3.5 μm, followed by deparaffinization, rehydration, and antigen retrieval by pressure-cooking (30 sec at 125°C and 10 sec at 90°C) in citrate buffer (10 mM, pH 6.0). After blocking for 1 h at room temperature with 5% bovine serum albumin, lung sections were incubated in primary antibodies overnight at 4 °C, followed by secondary antibody (1:250) incubation for 2h at room temperature. The following primary (1) and secondary (2) antibodies were used: (1) rat anti-Krt8 (170μg/ml; University of Iowa Hybridoma Bank, 1:200), rabbit anti-pro-SPC (1:200; Millipore, AB3786), goat anti-Pdpn (1:200; R&D Systems, AF3244), rabbit anti-SPC (1:150; Sigma-Aldrich, HPA010928), mouse anti-alphaSMA (1:1,000, Sigma-Aldrich, A5228), rabbit anti-Areg (1:50; LSBIO, LS-B13911), rabbit anti-Hbegf (1:200; Bioss Antibodies, bs-3576R), rabbit anti-Ki67 (1:200; Abcam, ab16667), mouse anti-CC10 (1:200; Santa Cruz, sc-365992), rabbit anti-Cst3 (1:100; Abcam, ab109508); (2) donkey anti-rabbit AlexaFluor568 (Invitrogen, A10042), donkey anti-rat AlexaFluor488 (Invitrogen, A21208), donkey anti-goat AlexaFluor647 (Invitrogen, A21447), goat anti-mouse AlexaFluor647 (Invitrogen, A21236). Images were acquired with an LSM 710 microscope (Zeiss).

### Microscopic image analysis - quantification

The fluorescence intensity of Krt8 expression in selected regions of immunofluorescence microscopy images was measured excluding airways using FIJI (ImageJ)^81^. For quantification of Krt8 expression in the human FFPE sections, and likewise, for the Hbegf and Areg quantification in the mouse sections, the mean overall fluorescence intensities were measured. For quantification of cell proliferation, cells were stained with Ki67 and Krt8 and counted manually for Ki67 positive cells.

### Lineage tracing experiments

SPC-CreERT2 (Sftpc^tm1(cre/ERT2,rtTA)Hap^) mice were crossed with Gt(ROSA)26Sortm4(ACTB-tdTomato,-EGFP)Luo mice. Sox2-CreERT2 mice were crossed with Ai14-tdTomato (Gt(ROSA)26Sor^tm14(CAG-tdTomato)Hze^). Four doses (SPC-CreERT2) or three doses (Sox2-CreERT2) of 0.25 mg/g body weight tamoxifen in 50 μl corn oil. A chase period of >21 days was used to ensure the absence of residual tamoxifen before injury. Bleomycin (1.5 U/kg) was delivered to mouse lungs via oral aspiration in 40 μL sterile PBS. Lungs were harvested at 10 days or 14 days following injury.

### Hypoxia/Hyperoxia + InfA infection model

Wild-type or bi-transgenic Sftpc^CreERT2^; Rosa26R^mTmG^ mice were exposed to 12% (hypoxia), 21% (room air) or 100% (hyperoxia) oxygen between postnatal days 0-4^82^. All mice were then exposed to room air until they were 8 weeks old. Sftpc^CreERT2^; Rosa26R^mTmG^ mice were administered tamoxifen (Sigma Aldrich, T5648) (0.25g/kg) or corn oil vehicle by single daily injections for four consecutive days^38^. On the seventh day, the mice were infected with influenza A virus (HKx31, H3N2) and lungs were harvested on post-infection day 14. Lungs were inflation fixed overnight in 10% neutral buffered formalin, embedded in paraffin, sectioned and stained with antibodies against pro-SPC (Seven Hills Bioreagents, Cincinnati, OH);

T1alpha (1:100; Syrian Hamster, clone 8.1.1, DSHB-Developmental Studies Hybridoma Bank at the University of Iowa) and Krt8/TROMA-I (DSHB-Developmental Studies Hybridoma Bank at the University of Iowa); Sections were incubated with fluorescently labeled secondary antibody and stained with 4’, 6-diamidino-2-phenylindole (DAPI). Slides were visualized with a Nikon E-800 fluorescence microscope (Nikon Instruments, Microvideo Instruments, Avon, MA). Images were captured with a SPOT-RT digital camera (Diagnostic Instruments, Sterling Heights, MI).

## Acknowledgements

We gratefully acknowledge the provision of human biomaterial and clinical data from the CPC-M bioArchive and its partners at the Asklepios Biobank Gauting, the Klinikum der Universität München and Ludwig-Maximilians-Universität München. The Krt8/TROMA-I monoclonal antibody, developed by Brulet, P. / Kemler, R. was obtained from the Developmental Studies Hybridoma Bank, created by the NICHD of the NIH and maintained at the University of Iowa, Department of Biology, Iowa City, IA, 52242. The beta6 integrin (Itgb6, clone 3G9) antibody was kindly provided by Prof. Dr. Dean Sheppard at the University of California San Francisco.

## Author contributions

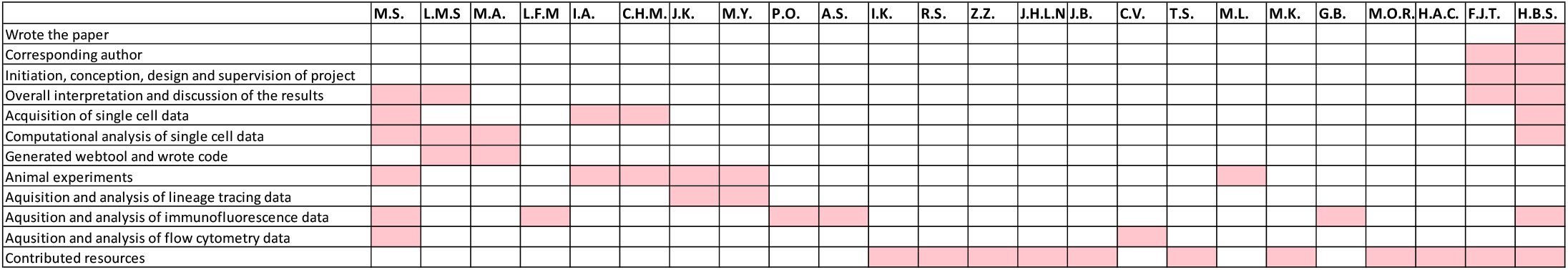

---

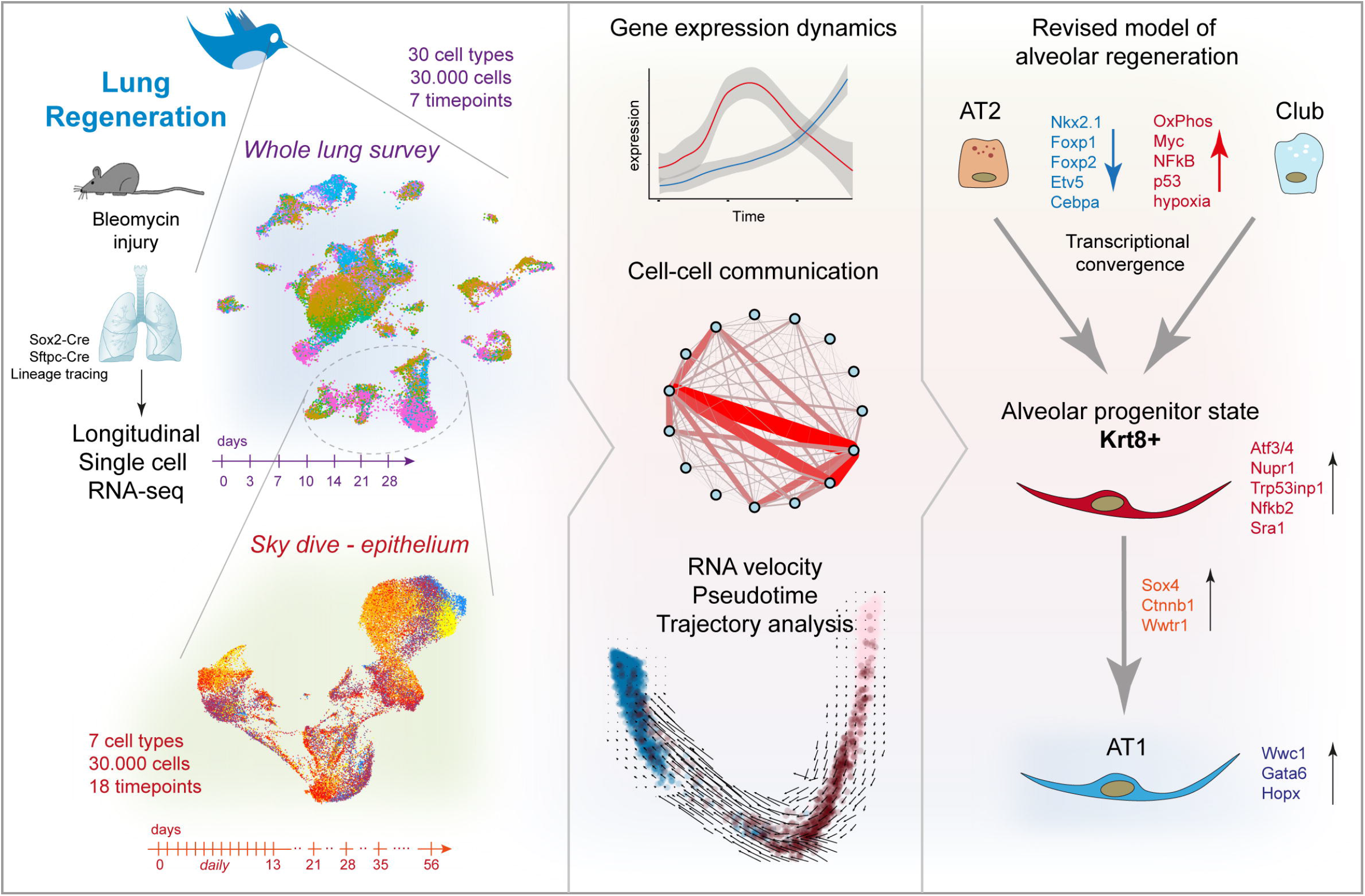

